# A genomewide association study for bristle number variation in *Drosophila melanogaster*

**DOI:** 10.64898/2026.08.17.745380

**Authors:** Katherine M. Hanson, Stuart J. Macdonald

**Author notes:** Corresponding author: (SJM).

## Abstract

Decades of research has uncovered a wealth of mechanistic information about the development of sensory bristles in *Drosophila melanogaster*. By studying large-effect, often loss-of-function mutations, many genes have been associated with bristle development, morphology, patterning, and number. Equally, the number of bristles present in certain areas of the fly cuticle is a classic quantitative trait, the genetic basis of which has been studied using a range of tools, from artificial selection to QTL (Quantitative Trait Locus) mapping. Such studies have often implicated well-understood bristle development genes as contributing to natural variation in bristle number. Here we contribute to the study of bristle number genetic variation in flies by executing a GWAS (genomewide association study). We generated whole genome sequences for 897 phenotyped male *D. melanogaster* individuals derived from a wild-derived, but lab-adapted outbred population, revealing - following quality control and filtering - over 780,000 variants with frequencies greater than 5%. Using these data we estimated the SNP (Single Nucleotide Polymorphism) heritability for ABN (abdominal bristle number) and SBN (sternopleural bristle number) as 0.28 and 0.35, respectively. These values indicate that our set of genotyped variants collectively explain a substantial fraction of the variance in phenotype in the mapping panel. Subsequently, genome scans revealed 1085 (ABN) and 211 (SBN) genomewide significant sites, and - due to extensive LD (Linkage Disequilibrium) in our panel - nearly all these sites are clustered into three locations; We find a GWAS hit for ABN in the middle of chromosome 3L, and hits for SBN at the tip of the X chromosome (where several prior mapping studies have resolved QTL for bristle number), and on 2L. Surveying existing studies that identified genes that control bristle number/development, we highlight several candidates that may segregate for causative, functional variants.

## Introduction

The bulk of traits of agricultural, medical, ecological and evolutionary significance exhibit quantitative variation among individuals in populations, and are controlled by diverse, genomewide sets of genetic variants, and an array of environmental factors. Work using various genetic mapping tools in a broad set of animal models, plant systems, and humans typically show that such complex traits are highly polygenic, with individual loci having only modest effects on phenotypic variation [e.g., 1, 2–6].

Tremendous insight into the biology of complex traits has been gained from both population-based GWAS (Genomewide Association Studies) and various forms of QTL (Quantitative Trait Locus) mapping in humans and non-human systems. The gain in knowledge and understanding has been most visible with human GWAS, where the genetic dissection of thousands of human diseases and disease-relevant traits in the last two decades has resulted in >625,000 significant variant-disease associations [7]. In turn, these associations have led to the identification of novel disease risk candidates [e.g., 8], enabled a range of functional work to elucidate the mechanisms of action of causative disease variants [9], and have aided in the prediction of disease risk [e.g., 10].

Genetic dissection of non-disease phenotypes in humans have also proven valuable in understanding the genetics and evolution of complex traits, and this has been particularly true of height. Since height is easy to measure with minimal error, GWAS with extremely large sample sizes have been executed [up to 5.4 million individuals; 11]. The power of these studies, coupled with the high heritability of the trait [12, 13], has enabled thousands of genotype-phenotype associations to be identified that collectively explain a significant fraction of the genetic variation for height [11, 14]. Other studies have also employed height as model trait to examine the contributions of common/rare variants to trait variation [15, 16], and to assess the effects of inbreeding [17] and selection [18, 19] on complex traits.

In the *Drosophila melanogaster* system, the number of bristles is a classic quantitative trait that - analagous to human height - has been widely-employed as a model complex trait [20–22]. Typically, two bristle traits are measured: Abdominal bristle number (or ABN) which is the number of bristles on a particular abdominal segment (sternite A5 in males and A6 in females), and sternopleural bristle number (or SBN) which is the sum of the number of bristles on the pair of lateral sternopleural plates.

Bristle number is a valuable model to enable the study of complex traits for at least three reasons. First, bristles are sensory organs, and their development, along with that of the entire fly peripheral nervous system, has been subject to a great deal of study [23]. As a result, many genes - *achaete*, *Delta*, *hairy*, *Notch*, *scute* to name a few - are known to be involved in bristle development, and they have served as useful targets to test whether they harbor allelic variation contributing to bristle number variation in populations [24–28]. Second, as a simple count-based trait, bristle number is easily and rapidly scored with limited error on intact, living (but anesthetized) animals. Furthermore, since the number and location of adult bristles are defined during development, the phenotypes are unaffected by adult age/condition. These factors enable straightforward experiments without the constraints that are often evident when studying behavioral or physiological traits. Third, bristle number shows relatively high heritability of 30-50% [29–33], and the strong genetic contribution to bristle number variation is further evidenced by the efficacy of artificial selection in shifting population means over a modest number of generations [33]. The ease of working with the trait and its high heritability has led to a number of studies seeking to genetically dissect bristle number variation [24, 26–28, 33–45], and to screen for mutations impacting the trait [46–48]. These studies have added to the understanding of the genetic variation contributing to bristle number, but have also enabled the development and validation of novel experimental approaches that have subsequently been deployed to genetically dissect other traits.

One approach that has yet to be employed to study bristle number variation in *D. melanogaster* is a GWAS. While studies have employed gene-centric association mapping - i.e., testing variants in/near a single locus for their effects on bristle number [e.g., 26, 44] - no study has attempted a true, unbiased genomewide experiment. Here we present such a study. We employ ∼900 phenotyped and whole-genome sequenced *D. melanogaster* males derived from a lab-adapted, but wild-derived population. A genome scan testing effects at >780,000 variants revealed >1,200 variants associated with phenotype, the vast majority of which reside in 3 clusters, and appear to implicate previously-known bristle number candidate genes.

## Materials and Methods

### Mapping population

The population we employed was founded by the progeny of several hundred *Drosophila melanogaster* male and female flies that were wild-caught at the Fenn Valley Vineyards (Fennville, Michigan, USA) in 2010 by Dr. Ian Dworkin’s research group. This FVW population was subsequently maintained in the Dworkin lab [see 49 for additional detail]. In 2014 we received several bottles of embryos/larvae derived from the FVW population, and established a copy of the population in our lab. This was maintained for several years by provisioning the cage weekly with bottles (6-oz, Fisher Scientific, AS115) of fresh cornmeal-yeast-molasses media (S1 Text).

### Experimental fly rearing and phenotyping

Embryos were collected from our FVW population cage as follows. We added to the cage several 100-mm petri dishes containing apple juice agar (S2 Text) supplemented with a small amount of live yeast paste. After 24 hours, dishes were removed from the cage, and eggs were removed from the agar surface, rinsed, and resuspended in 1X PBS (phosphate-buffered saline solution). Subsequently, we moved 12-μl of resuspended eggs to each of ∼100 standard, narrow fly vials (Fisher Scientific, AS515) each containing 10-ml of cornmeal-yeast-molasses media (S1 Text). Collecting eggs in this way yields a fairly consistent larval density over culture vials [50]. More detail on the egg collection procedure is provided in S3 Text.

Eggs were reared into experimental adults in an incubator set to 25°C, 50% relative humidity, with a 12 hour light : 12 hour dark cycle. Adults were phenotyped 12-24 days following egg collection, and to maintain reasonable vial conditions, adults were periodically transferred to fresh media vials during this period. For a maximum of 12 adult males per source vial we counted/recorded abdominal and sternopleural bristle number (ABN and SBN, respectively). ABN is number of bristles on the most-posterior ventral sternite, which in male *D. melanogaster* corresponds to segment five [see Figure 46b in 51], while SBN is the sum of the number of bristles on the right and left sternopleural plates [see Figure 31 in 51]. See S1 Table for the raw bristle count data. ABN and SBN are the “standard” bristle phenotypes collected by researchers over at least the last 30 years [for instance, 33]. Phenotyped animals were deposited individually in wells of a series of 96-well plates (500-μl well, round bottom; Axygen, P-96-450R-C) that were held on ice during phenotyping, and subsequently frozen at −20°C until DNA isolation.

### DNA isolation, library construction and sequencing

DNA was isolated from 960 animals in a series of 96-well plates using an adaptation of the Qiagen Puregene Cell Kit (catalog number 158046; see S4 Text), and all samples were quantified using the Qubit dsDNA Broad Range kit (Invitrogen, Q32850). Libraries were prepared from this DNA using a custom, in-house Tn5 transposase-based method similar to the procedures outlined in Picelli et al. [52] and Hemmer et al. [53]. Briefly, 8-ng of genomic DNA was tagmented via Tn5 transposase, reactions were inactivated via the addition of SDS, then samples were amplified and barcoded via PCR using a series of unique dual indexed primers (see S5 Text S5 and S2 Table for details on the protocol and primers). Following quantification of each individual library (Qubit dsDNA High Sensitivity kit; Invitrogen, Q32854), libraries were mixed by concentration into 12 pools, ignoring a subset of samples with low concentrations. Subsequently, each pool was cleaned up via a 0.8X AMPure XP bead cleanup (Beckman Coulter, A63881; S6 Text), size-selected to retain 400-600 bp fragments using a BluePippin instrument with a 2% agarose gel cassette (Sage Science, BDF2010), and cleaned up again via a 2X AMPure XP bead cleanup (S6 Text). Finally, the 12 pools were further mixed in groups of three for sequencing. Each of the final four pools contained 220-231 uniquely-indexed samples (for a total of 897 sequenced samples), had average fragment sizes of 588-598 bp (S1 Fig), and were sequenced on separate lanes of an Illumina NovaSeq 6000 S4 flowcell resulting in PE150 reads (Genewiz). On average we generated over 14 million raw read pairs per sample (mean = 14.1 million, standard deviation = 4.33 million).

### Read alignment, variant calling and variant filtering

Raw reads from each of the 897 sequenced male flies were mapped to Release 6 (dm6) of the *D. melanogaster* reference genome using bwa mem [version 0.7.17; 54], and duplicates were marked using the Picard toolkit [version 3.3; Broad 55]. On average, 94.37% of the reads aligned for each sample, with mean coverage of 12.5X for the X chromosome, and 22.6X for chromosomes 2 and 3. Given the relatively high level of variation in the *D. melanogaster* genome we additionally re-aligned bam files using the indel realigner from GATK [version 3.2-2; 56]. We initially identified 2,728,067 variants in the re-aligned bam files using bcftools [version 1.21; 57]. Then we used a combination of GATK [version 4.6.1.0; 56] and vcftools [verson 0.1.16; 58] to filter sites based on quality score, read depth, and to keep only biallelic sites with a minor allele frequency of at least 5%. This resulted in 1,596,977 variants. This set of variants was then pruned to remove sites in high linkage disequilibrium using PLINK [version 1.90b6.21; 59] with a window size of 50 variants, a step size of 10 variants, and an *r*^2^ threshold of 0.95. This pruning left 786,716 variants, which were tested for association with both bristle phenotypes in the 897 individuals.

### Genomewide scan for variants potentially associated with bristle number

We used GEMMA [version 0.98.5; 60] to both generate a centered relatedness matrix using the set of 786,716 variants, and to execute a GWAS for each of our bristle phenotypes using a linear mixed modeling framework that accounts for population stratification and relatedness among individuals. Variant-by-variant Wald test *P*-values for the two GWAS are provided in S3, S4, and S5 Tables. Significant GWAS associations were identified by adjusting each phenotype-specific set of *P*-values for multiple tests using the Benjamini and Hochberg [61] method using the p.adjust function in R [R Core62], and considering significant hits as those tests with adjusted *P*-values < 0.05. We acknowledge this threshold is not as stringent as a Bonferroni-corrected threshold (in our case 0.05/786,716 = 6.4 × 10^−8^) often used in GWAS. However, given the plethora of candidate genes that have been implicated in bristle number variation by prior work (see below), we felt that detecting even “suggestive” associations - particularly those in/near *a priori* candidate genes - may have utility.

### Calculating sites in Linkage Disequilibrium (LD) with lead variants

Within each of the 3 major GWAS hit regions (see below), we identified the most significant variant (the “lead variant”). Then we used PLINK [59] to generate an LD report, calculating LD (R^2^) between the lead variant and all variants within 3-Mb of it (see S6 Table and S2 Fig).

### Bristle number candidate genes

We generated a list of 432 bristle number candidate genes, nominating candidates in three ways. First, we used FlyBase vocabulary searches [release FB2021_06; 63] to extract all genes associated with the following terms: chaeta development (GO:0022416), lateral inhibition (GO:0046331), mechanosensory chaeta (FBbt:00005181), peripheral nervous system development (GO:0007422), proneural cluster (FBbt:00001135), and regulation of neurogenesis (GO:0050767). Second, we marked all genes that showed effects on abdominal or sternopleural bristle number following P-element mutagenesis, extracting this information from Table 2 of Norga et al. [48]. Third, we checked 22 publications that described plausible candidate genes: two bristle number quantitative genetics review articles [20, 21], four bristle number QTL mapping papers [33, 38, 40, 43], 15 bristle number genetic mapping studies that included functional tests for specific genes [22, 24, 26–28, 34–37, 39, 41, 42, 46, 64, 65], and one work that identified genes on the third chromosome that impact peripheral nervous system development [47], since such genes can impact bristle patterning. Details on the 432 candidates, and how they were implicated, is presented in S7 Table. Candidate genes within 20-Kb of a variant with a Benjamini-Hochberg-adjusted *P*-value < 0.05 are given in S8 Table.

### Data and software availability

Experimental protocols, sequences of custom sequencing library indexing oligos, bristle number phenotypes, association statistics, candidate genes, R code to reproduce figures from supplementary tables, and so on, are available as supplementary files. Raw sequencing data is available at the NCBI sequence read archive (SRA) under BioProject Accession PRJNA1506691, and all analytical code is available on GitHub (https://github.com/Hanson19/Bristle_GWAS/).

## Results and Discussion

### Observed bristle number variation is consistent with prior studies

The population used for the present study was initially founded with several hundred wild-caught *D. melanogaster* flies, and maintained in the lab for a number of years prior to sampling experimental animals. Variation in both abominal and sternopleural bristle number (ABN and SBN, respectively) appears approximately normally-distributed (Figs 1A and 1B), as has been observed in other outbred samples of flies [e.g., 66]. The level of variation we see in both phenotypes - ABN = 17.8 ± 2.03, SBN = 18.1 ± 1.79 (mean ± 1-SD) - is also not dissimilar to that seen in prior studies of sets of outbred animals and collections of wild-derived inbred strains [28, 45, 66, 67]. Finally, we note that there is a low, but significant positive correlation between ABN and SBN across animals (*r* = 0.08, *P* = 0.015; Fig 1C). This likely reflects the modest genetic correlation between these traits [37, 46], which is also evidenced by studies showing that some mutations can have pleiotropic effects on both traits [e.g., 46, 48], and is - at least in part - responsible for the correlated response of SBN to artificial selection on ABN [33, 68].

**Fig 1.**
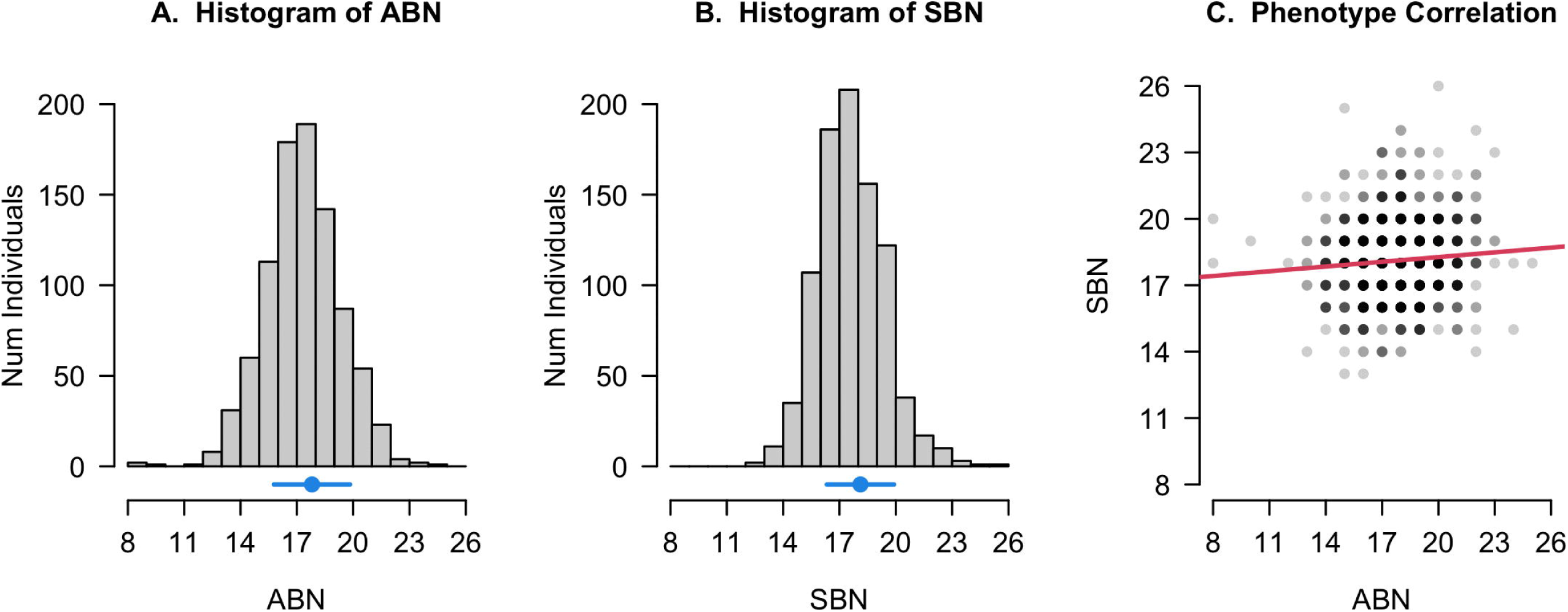
Bristle number variation. (A) and (B) Distributions of Abdominal Bristle Number, ABN, and Sternopleural Bristle number, SBN, among 897 phenotyped individuals. The phenotype means (± 1-SD) are shown in blue above each *x*-axis (ABN, 17.8 ± 2.03; SBN, 18.1 ± 1.79). (C) Limited correlation between the two phenotypes (Pearson’s *r* = 0.08, *P* = 0.015). Points are darker when more individuals share the same bristle counts.

### Estimates of SNP heritability for bristle number from several hundred sequenced individuals

We phenotyped ABN and SBN for 897 outbred *D. melanogaster* males, and generated whole genome sequencing data from each animal. Following a series of filtering steps (see Materials & Methods) a set of 786,716 common variants were available for testing. These data enable an estimate of so-called “SNP heritability” [see 69], the fraction of the variance in phenotype collectively explained by the genotyped variants. GEMMA [60] estimated the SNP heritabilities of ABN as 0.28 (standard error = 0.054) and SBN as 0.35 (standard error = 0.056). These values are lower than previous estimates of bristle number heritability [e.g., from artificial selection experiments such as 33]. However, we note that our sample size (N=897) is modest, we purged variants with minor allele frequencies below 5%, and not all causative variants may be strong LD with the one of the set of genotyped variants, all of which are likely to underestimate heritability.

### A GWAS implicates three genomic regions impacting bristle number

We used GEMMA to execute a GWAS for each of our target traits, employing a linear mixed modeling framework [60]. Fig 2 shows the results of these genome scans for each trait. We elected to treat variants as being significantly associated with a trait if the trait-specific test had a Benjamini-Hochberg adjusted *P*-value below 0.05 (red points in Fig 2). This yielded 1296 associated variants, 1085 for ABN (Fig 2A), and 211 for SBN (Fig 2B). No variant was significant for both traits, and the bulk of significant sites (97.8%) are located in three clusters: On chromosome 3L for ABN (1058/1085 sites), and on the X (85/211 sites) and on 2L for SBN (124/211 sites). We note that a harsher, per-trait Bonferroni correction (i.e., 0.05/786,716 = 6.4 × 10^−8^) implicates 72 variants for ABN, all of which reside in the 3L cluster, and 1 variant for SBN, which is the lead variant (i.e., the most significant variant) for the SBN 2L cluster.

**Fig 2.**
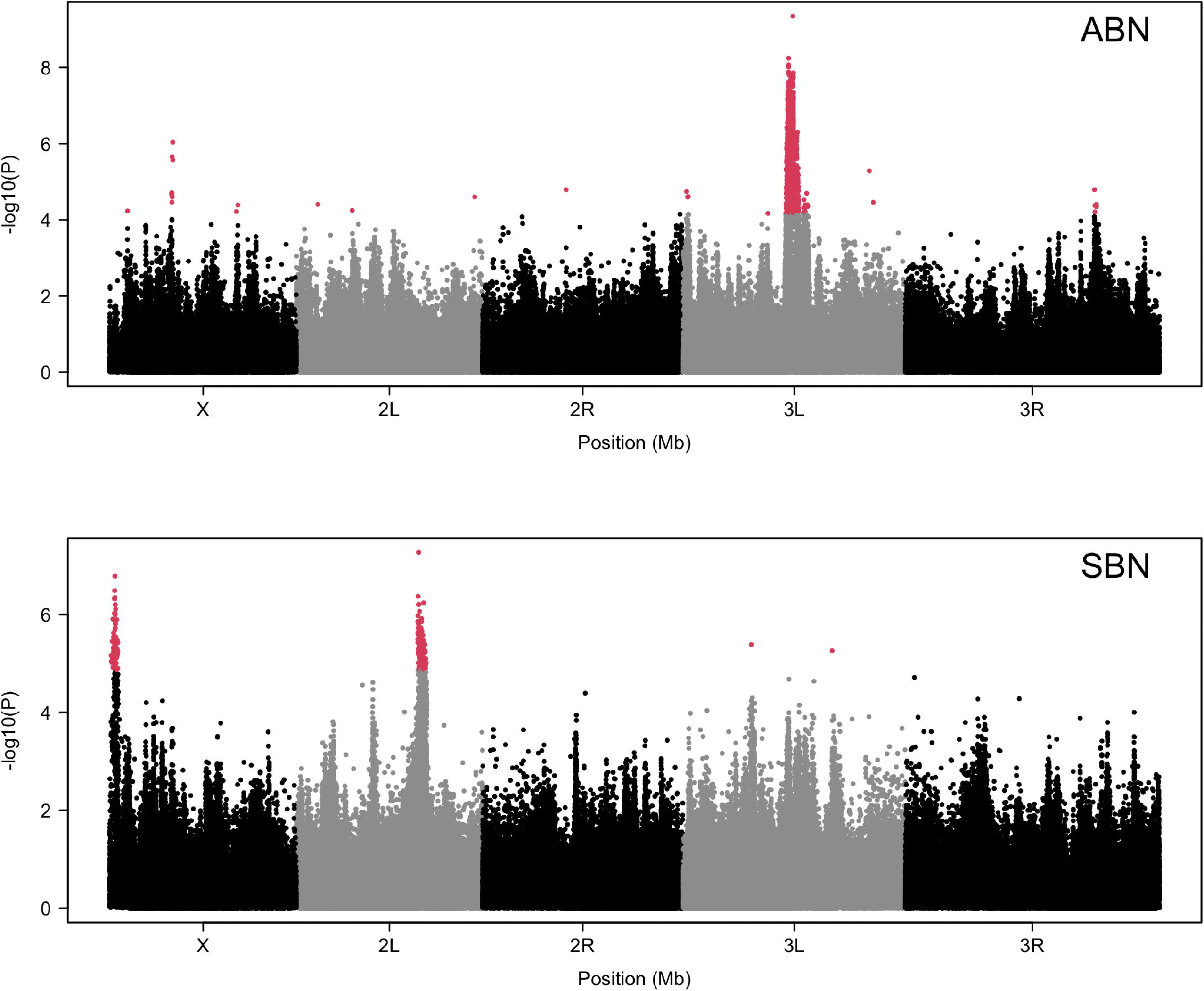
Manhattan plots for GWAS on two bristle traits in males. Each point is the physical location of one of the 783,878 variants tested along the 5 major *D. melanogaster* chromosome arms. The −log_10_(*P*-values) result from Wald tests for association with variation in abdominal bristle number (ABN, top panel) and sternopleural bristle number (SBN, bottom panel). Variants are colored in red if the Benjamini-Hochberg adjusted *P*-value is less than 0.05. There are 1,296 such sites (1,085 for ABN, 211 for SBN), and the vast majority (1,267, 97.8%) are clustered in 3 regions.

These three regions are the most likely to contribute to bristle number variation in our study. To explore them further we generated localized Manhattan plots (Figs 3 and 4). These plots illustrate the fairly extensive LD present in our sample of flies; strong LD (R^2^ > 0.8) between the lead variant and other variants in the cluster is evident at distances up to 0.5-Mb (see also S2 Fig). The lengths of the intervals over which we see strong LD are much greater than seen in previous studies using wild-caught animals or wild-derived inbred strains, where LD decays to a low level within a few kilobases [27, 44, 70]. The extended LD plausibly results from a combination of the founding of our population using just a few hundred genotypes, and its maintenance over several years (and many generations) in a laboratory population cage, likely limiting the effective population size. While impacting our resolution (see below) the extended level of LD may have actually increased our ability to find plausible associations with phenotype in our study; A GWAS sample size of 900 provides only limited power to detect sites that make only modest contributions to phenotypic variance [71], and with higher LD one is more likely to capture effects of causal variants in with a given set of marker variants.

**Figure 3.**
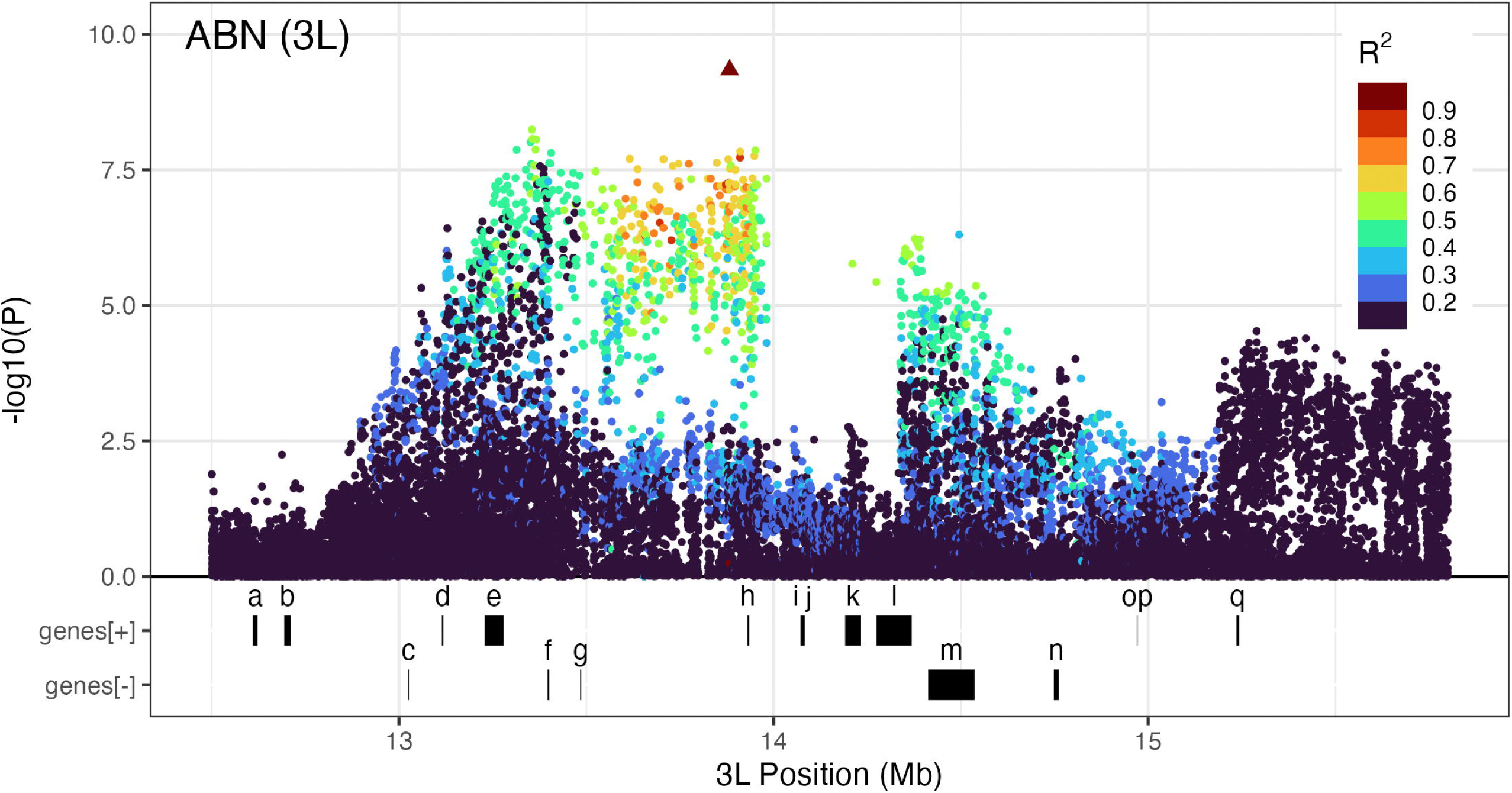
Zoomed-in Manhattan plot for chromosome 3L ABN GWAS hit. Physical locations of variants are plotted against the −log_10_(*P*-values) resulting from Wald tests for association with abdominal bristle number (ABN). The “lead” variant in the region is marked with a triangle. All other variants (solid circles) are color-coded based on their linkage disequilibrium (LD, as measured by *R*^2^) with the lead variant. At the bottom of the plot are the locations of bristle number candidate genes on the plus and minus strands. Letters correspond to the following genes: (a) *caup*, (b) *mirr*, (c) *Zmynd10*, (d) *trn*, (e) *caps*, (f) *sens*, (g) *Abp1*, (h) *DCTN1-p150*, (i) *Pex1*, (j) *btl*, (k) *nuf*, (l) *fz*, (m) *bbg*, (n) *Trl*, (o) *Tom*, (p) *Brd*, (q) *Tollo*. Note that the locations of the pairs of genes marked with i and j, and with o and p, are too close to be distinguished at the resolution of this figure.

**Figure 4.**
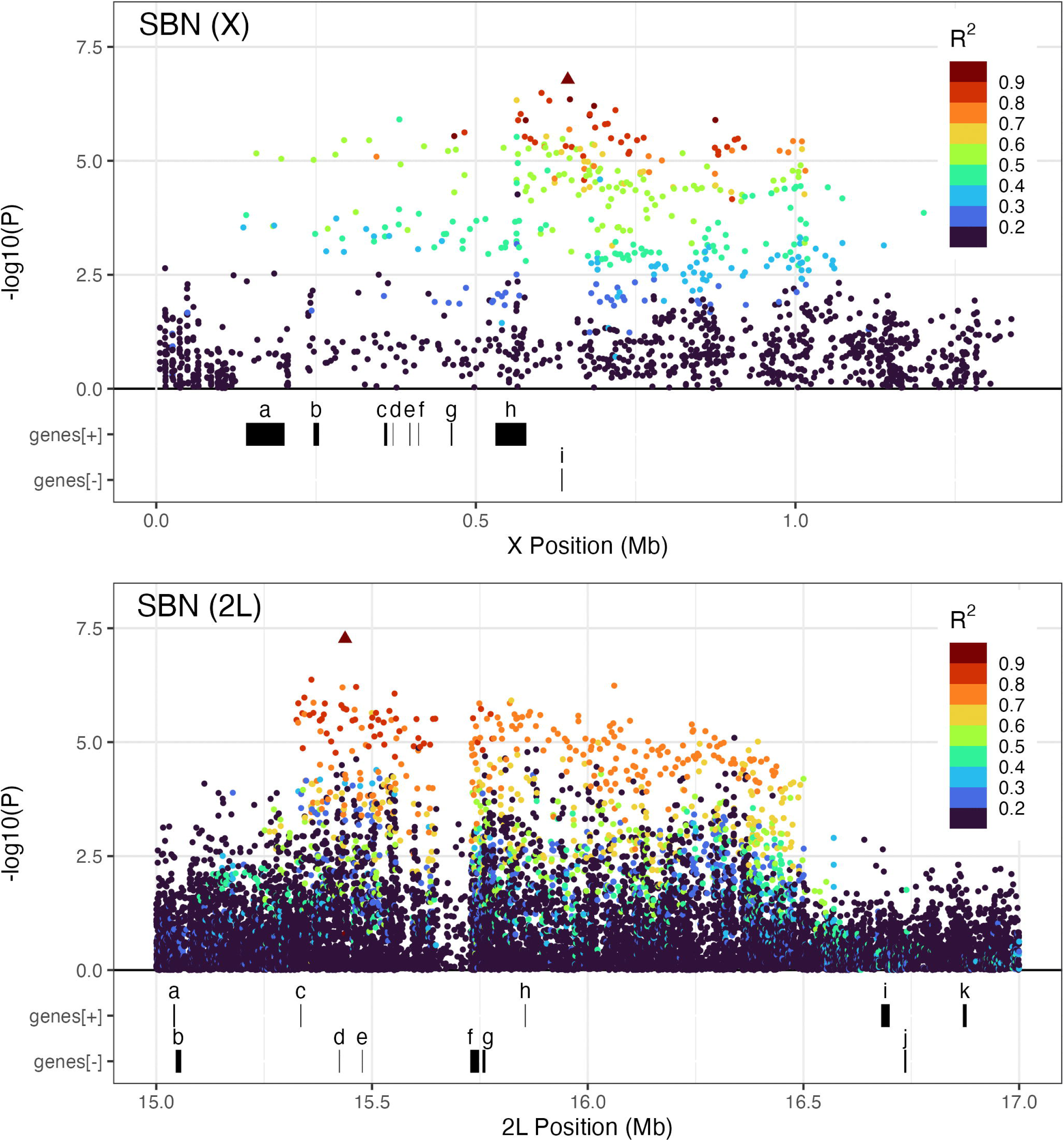
Zoomed-in Manhattan plot for the two major SBN GWAS hits. The top plot shows the hit on the X, and the bottom shows the hit on chromosome 2L. In each plot the physical locations of variants are plotted against the −log_10_(*P*-values) coming from Wald association tests with sternopleural bristle number (SBN). The “lead” variant in each region is represented with a triangle, while all other variants (solid circles) are color-coded based on their linkage disequilibrium (LD measured by *R*^2^) with the lead variant. Bristle number candidate gene locations on the plus and minus strands are at the bottom of each plot. For the X-chromosome hit letters correspond to the following genes: (a) *tyn*, (b) *G9a*, (c) *y*, (d) *ac*, (e) *sc*, (f) *l(1)sc*, (g) *ase*, (h) *Appl*, (i) *Dredd*. For the hit on 2L letters correspond to: (a) *Su(H)*, (b) *ck*, (c) *esg*, (d) *wor*, (e) *sna*, (f) *CycE*, (g) *Gli*, (h) *Tektin-A*, (i) *grp*, (j) *glu*, (k) *lncRNA:CR31781*.

### Connecting GWAS hits to *a priori* bristle number candidate genes

The development of bristles, and the genetic basis of bristle number, have been studied in *Drosophila* for decades. As such, many genes are known to impact bristles, and these are plausible candidates to harbor functional genetic variants leading to variation in bristle number in the present study. We identified 432 such candidates from a range of sources (see Materials & Methods) and sought to use these to inform our association results.

The localized Manhattan plots depicting the three clusters of GWAS hits each highlight the series of bristle number candidate genes residing in the intervals (Figs 3 and 4). Additionally, we marked those candidate genes that are within 20-kb of any GWAS hit, including those hits not within the three primary clusters (S8 Table). We acknowledge that 20-kb is an arbitrary distance threshold. An alternative approach may have been to select those genes with hits within the transcribed region. But given the extensive LD in our mapping population (above), and given that most variants contributing to complex trait variation are expected to be in non-coding regions and have regulatory effects [72], that strategy might exclude plausible causatives. Conversely, if we were to use a much longer distance threshold we would include a much larger number of candidates, and would be more likely to include genes that do not actually impact bristle number variation in our population.

Seventeen bristle number candidate genes are within the cluster of ABN hits on chromosome 3L, and nine of these genes are within 20-kb of a hit: *trn*, *caps*, *sens*, *Abp1*, *DCTN1-p150*, *nuf*, *fz*, *bbg*, and *Tollo* (S8 Table). *DCTN1-p150* is the closest candidate to the lead variant (Fig 3, gene “h” below the x-axis), although is only implicated based on its associated with the GO term GO:0050767 (regulation of neurogenesis). Among the other eight genes, both *trn* (gene “d”) and *sens* (gene “f”) have been shown to be involved in development of the peripheral nervous system in a P-element screen [47]. Furthermore, *trn* has been implicated by previous genetic mapping studies, being near to QTL peaks mapped for SBN [43] and for both bristle traits [40].

All nine of the bristle number candidates within the SBN hit interval on the X chromosome (Fig 4, top panel), are also within 20-kb of a hit (S8 Table). The gene closest to the lead variant is *Dredd* (gene “i” below the x-axis in Fig 4, top panel), which is implicated as a bristle number candidate via the GO term GO:0007422 (peripheral nervous system development). However, arguably the strongest candidates in this interval are *ac* (gene “d”) and *sc* (gene “e”); variation at these genes has been previously associated with bristle number [28, 35], and studies have also mapped QTL to the region containing them [33, 38, 40, 43].

Six of the eleven genes within the SBN hit interval on chromosome 2L are within 20-kb of a GWAS hit (Fig 4, bottom panel; S8 Table): *esg*, *wor*, *sna*, *CycE*, *Gli*, and *Tektin-A*. Five of these are implicated as bristle candidates solely via GO terms (S7 Table), while *esg* (gene “c” below the x-axis in Fig 4, bottom panel) emerged from a screen to identify insertion mutations impacting bristle number [48].

Finally, we implicated five additional candidates that reside within 20-kb of one or more hits outside of the three GWAS hit clusters, all of which were hits for ABN (S8 Table). The gene *sws* - included as a candidate since it was identified in a screen for mutants impacting bristle number [48] - is within 20-kb of one of a set of 8 ABN hits on the X (Fig 2, top panel). The *Root* gene - marked with the FlyBase vocabulary term “proneural cluster” (FBbt:00001135) - is within 20-kb of one of the set of 5 ABN hits on 3R (Fig 2, top panel). While the remaining three genes - *exd*, *lola*, and *psq* - are within 20-kb of isolated GWAS hits.

### Summary and caveats

Our GWAS results support the role of a handful of genes that have been previously implicated in the development of bristles, and/or have been associated with variation in bristle number via a range of genetic mapping studies and screens. This being said, as with any unbiased genetic mapping study, confirmation of the involvement of these genes in contributing to phenotype via the use of targeted functional genetic tests is critical. Clearly, while we focus on *a priori* bristle candidates, our study does not rule out the possibility that none of those genes we highlight above harbor functional, phenotypically-relevant variants in our base population; Other genes not previously connected to bristle number may be causative. Studies with greater power and better resolution than we present - likely an association study using a much larger set of wild-caught animals, where the population exhibits shorter tracts of LD - would enable improved dissection of the trait and the genes involved, and a less speculative route to functional validation. With the costs of sequencing continuing to fall, it is plausible that such studies are achievable in the near future.

## Acknowledgements

We thank Kristen Cloud-Richardson for molecular biology assistance, and Brian Sanderson for computational support. Analytical work was supported by infrastructure provided through the Data Science Core of the Kansas INBRE project, the KU Center for Genomics, and the KU Center for Research Computing cluster. We also thank Brittny Smith in the KU Genome Sequencing Core for assistance with sequencing library quality control and cleanup.

## Supporting Information Captions

**S1 Text.** Ingredients and brief protocol for Macdonald lab cornmeal-yeast-molasses fly media.

**S2 Text.** Ingredients and brief protocol for Macdonald lab apple juice agar plates.

**S3 Text.** Macdonald lab egg collection protocol.

**S4 Text.** Macdonald lab 96-well plate DNA isolation protocol.

**S5 Text.** Custom, in-house Tn5-based library preparation protocol.

**S6 Text.** Library bead-cleanup protocols.

**S7 Text.** R code enabling reproduction of figures from supplementary tables.

**S1 Table. Phenotype data.** Bristle count data for the 897 phenotyped *Drosophila melanogaster* males. ABN = Abdominal Bristle Number. SBN = Sternopleural Bristle Number.

**S2 Table.** Unique dual indexing oligos used for sequencing library amplification and barcoding.

**S3 Table. GWAS *P*-values.** Results from the 786,716 association tests (file 1 of 3). Column “chr” = Chromosome arm (X, 2L, 2R, 3L, 3R, 4, Y). Column “pos_R6” = Variant location in the *Drosophila melanogaster* reference genome (Release 6 coordinates). Columns “p_wald_abn” and “p_wald_sbn” = Raw variant-by-variant *P*-values resulting from Wald tests in GEMMA for abdominal bristle number (abn) and sternopleural bristle number (sbn).

**S4 Table. GWAS *P*-values.** Results from the 786,716 association tests (file 2 of 3). Column “chr” = Chromosome arm (X, 2L, 2R, 3L, 3R, 4, Y). Column “pos_R6” = Variant location in the *Drosophila melanogaster* reference genome (Release 6 coordinates). Columns “p_wald_abn” and “p_wald_sbn” = Raw variant-by-variant *P*-values resulting from Wald tests in GEMMA for abdominal bristle number (abn) and sternopleural bristle number (sbn).

**S5 Table. GWAS *P*-values.** Results from the 786,716 association tests (file 3 of 3). Column “chr” = Chromosome arm (X, 2L, 2R, 3L, 3R, 4, Y). Column “pos_R6” = Variant location in the *Drosophila melanogaster* reference genome (Release 6 coordinates). Columns “p_wald_abn” and “p_wald_sbn” = Raw variant-by-variant *P*-values resulting from Wald tests in GEMMA for abdominal bristle number (abn) and sternopleural bristle number (sbn).

**S6 Table. LD estimates.** Estimates of Linkage Disequilibrium (LD) between the lead variant at each of the 3 major GWAS hit regions and other variants within 3-Mb. Column “pheno” = The phenotype underlying the GWAS hit (ABN, SBN). Column “chr” = The chromosome arm on which the hit resides (chrX, chr2L, chr3L). Column “lead” = Location of the lead variant (i.e. the variant with the lowest *P*-value in the region) for each of the 3 GWAS hit regions in the *Drosophila melanogaster* reference genome (Release 6 coordinates). Column “pos_R6” = Variant location in the *Drosophila melanogaster* reference genome (Release 6). Column “R2” = PLINK-calculated LD (R^2^) between the two variants.

**S7 Table. Bristle candidate genes.** A list of 432 bristle number candidate genes derived from the literature and other prior work. Column “Gene_Table_Order” = A 1:432 vector ordering the table by the value in the “Gene_Symbol” column. Columns “FBgn”, “Gene_Symbol”, “Gene_Name”, and “CG_CR” = Names/IDs for the genes. Columns “Chr_Arm”, “Pos_Min_R6”, “Pos_Max_R6”, and “Strand” = Location information for the genes. Column “PMID_List” = A forward slash separated list of the PubMed IDs that implicated the gene (see main text for the list of 22 publications screened). Or “NA” if no publication implicated the gene. Column “How_Gene_Is_Implicated_Based_On_PMIDs” = Can take 1 of 5 levels. (1) only_suggested_in_review = Gene only presented in a review paper. (2) implicated_via_qtl = Gene is stated as being implicated by a mapped QTL (+ may also be in a review). (3) test_at_gene_PNS = Gene implicated via a gene-specific test for a PNS phenotype (+ may also be implicated by a QTL and/or in a review). (4) test_at_gene_bristle = Gene implicated via a gene-specific test for a bristle phenotype (+ may also be implicated via a test on a PNS phenotype and/or implicated by QTL and/or in a review). Column “Norga_Hit” = Takes a value of “yes” or “no” based on Table 2 of Norga et al. (2003, PMID: 12932322) which used P-element insertions to identify genes with effects on bristle number. Column “GO_Term_List” = A forward slash separated list of FlyBase vocabulary terms that implicated the gene. Or “NA” if none did.

**S8 Table.** Set of bristle number candidate genes that are within 20-kb of a GWAS association hit (adjusted *P*-value < 0.05). Genes noted as “in cluster” are present within one of the 3 clusters of hits on 3L (ABN), X (SBN) and 2L (SBN).

**S1 Fig.** Fragment size distributions of the 4 sequenced library pools. Run on a high sensitivity D1000 ScreenTape on a Agilent Technologies TapeStation. Images provided by Genewiz.

**S2 Fig.** Linkage Disequilibrium (LD) at each cluster of GWAS hits. Plots of LD (in R^2^) between the lead variant (that with the lowest *P*-value) at each location and all other variants within 3-Mb of it.

