## Supplementary material for "A genomewide association study for bristle number variation in *Drosophila melanogaster*": S1_Fig.pdf

**S1 Fig.** Size distributions of the 4 sequenced library pools. Run on a high sensitivity D1000 ScreenTape on a Agilent Technologies TapeStation. Images provided by Genewiz.

**B1: SM01-SJM-FVW-POOL-A2**

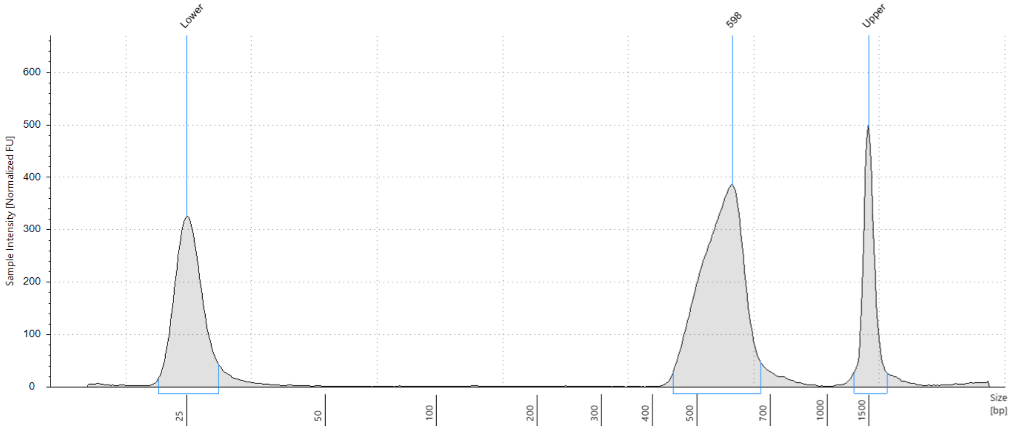


**C1: SM02-SJM-FVW-POOL-B2**

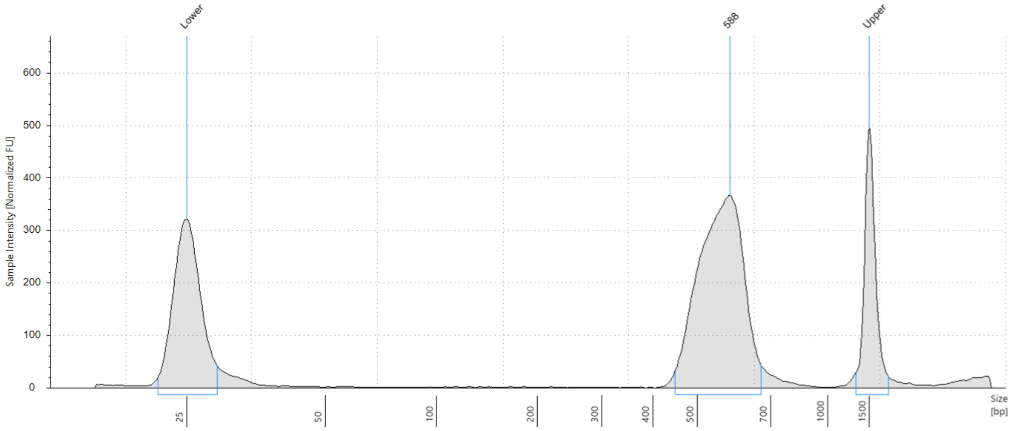

**S1 Fig.** Contd.

**D1: SM03-SJM-FVW-POOL-C2**

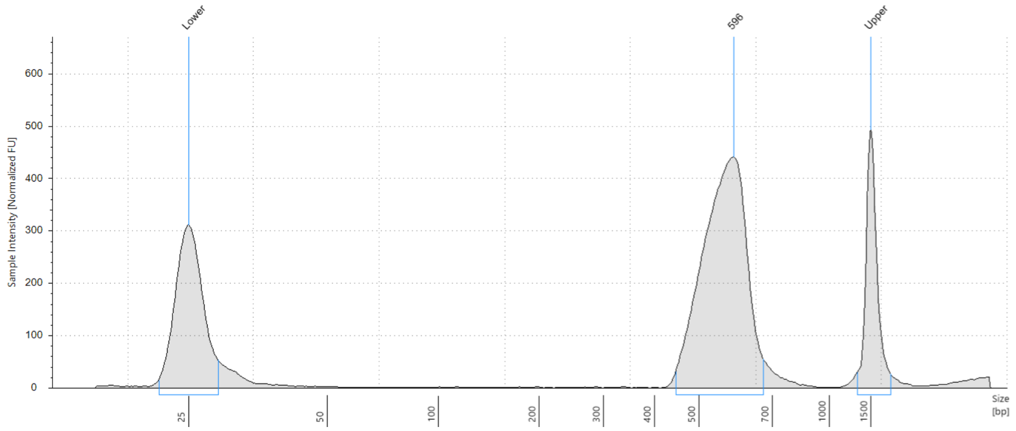


**E1: SM04-SJM-FVW-POOL-D2**

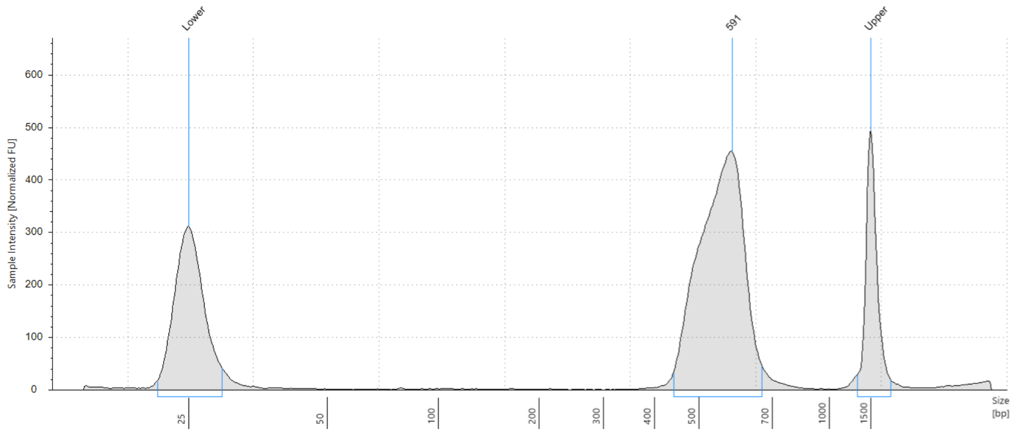
