## Supplementary material for "A genomewide association study for bristle number variation in *Drosophila melanogaster*": S1_Table.pdf

**S1 Table. Phenotype data.** Bristle count data for the 897 phenotyped *Drosophila melanogaster* males (ABN, Abdominal Bristle Number; SBN, Sternopleural Bristle Number).

| Sample_ID | ABN | SBN |
|---|---|---|
| FVWm_15_19_L02_A01 | 15 | 19 |
| FVWm_24_18_L02_B01 | 24 | 18 |
| FVWm_17_20_L02_C01 | 17 | 20 |
| FVWm_19_17_L02_F01 | 19 | 17 |
| FVWm_16_20_L02_G01 | 16 | 20 |
| FVWm_14_15_L02_H01 | 14 | 15 |
| FVWm_19_19_L02_D02 | 19 | 19 |
| FVWm_20_19_L02_E02 | 20 | 19 |
| FVWm_16_17_L02_F02 | 16 | 17 |
| FVWm_16_18_L02_G02 | 16 | 18 |
| FVWm_16_21_L02_B03 | 16 | 21 |
| FVWm_20_17_L02_C03 | 20 | 17 |
| FVWm_18_19_L02_D03 | 18 | 19 |
| FVWm_16_18_L02_E03 | 16 | 18 |
| FVWm_20_20_L02_F03 | 20 | 20 |
| FVWm_17_16_L02_G03 | 17 | 16 |
| FVWm_19_16_L02_A04 | 19 | 16 |
| FVWm_17_20_L02_B04 | 17 | 20 |
| FVWm_16_20_L02_C04 | 16 | 20 |
| FVWm_18_19_L02_D04 | 18 | 19 |
| FVWm_17_16_L02_E04 | 17 | 16 |
| FVWm_17_19_L02_F04 | 17 | 19 |
| FVWm_19_20_L02_G04 | 19 | 20 |
| FVWm_17_16_L02_H04 | 17 | 16 |
| FVWm_18_16_L02_D05 | 18 | 16 |
| FVWm_19_17_L02_E05 | 19 | 17 |
| FVWm_16_17_L02_G05 | 16 | 17 |
| FVWm_20_18_L02_A06 | 20 | 18 |
| FVWm_17_17_L02_D06 | 17 | 17 |
| FVWm_17_18_L02_E06 | 17 | 18 |
| FVWm_15_19_L02_G06 | 15 | 19 |
| FVWm_19_17_L02_A07 | 19 | 17 |
| FVWm_19_17_L02_B07 | 19 | 17 |
| FVWm_18_19_L02_C07 | 18 | 19 |
| FVWm_19_19_L02_D07 | 19 | 19 |
| FVWm_18_21_L02_E07 | 18 | 21 |
| FVWm_16_22_L02_F07 | 16 | 22 |
| FVWm_21_19_L02_H07 | 21 | 19 |
| FVWm_16_19_L02_B08 | 16 | 19 |
| FVWm_18_18_L02_C08 | 18 | 18 |
| FVWm_17_20_L02_D08 | 17 | 20 |
| FVWm_21_18_L02_E08 | 21 | 18 |
| FVWm_17_17_L02_G08 | 17 | 17 |
| FVWm_15_20_L02_H08 | 15 | 20 |
| FVWm_18_22_L02_C09 | 18 | 22 |
| FVWm_18_19_L02_E09 | 18 | 19 |
| FVWm_17_17_L02_F09 | 17 | 17 |
| FVWm_18_18_L02_G09 | 18 | 18 |
| FVWm_18_17_L02_H09 | 18 | 17 |
| FVWm_19_18_L02_A10 | 19 | 18 |
| FVWm_18_17_L02_B10 | 18 | 17 |
| FVWm_18_18_L02_D10 | 18 | 18 |
| FVWm_15_19_L02_E10 | 15 | 19 |
| FVWm_21_18_L02_F10 | 21 | 18 |

| Sample_ID | ABN | SBN |
|---|---|---|
| FVWm_19_16_L06_G12 | 19 | 16 |
| FVWm_21_19_L06_H12 | 21 | 19 |
| FVWm_19_15_L07_A01 | 19 | 15 |
| FVWm_17_19_L07_B01 | 17 | 19 |
| FVWm_15_15_L07_D01 | 15 | 15 |
| FVWm_18_17_L07_E01 | 18 | 17 |
| FVWm_19_15_L07_F01 | 19 | 15 |
| FVWm_14_17_L07_G01 | 14 | 17 |
| FVWm_14_18_L07_A02 | 14 | 18 |
| FVWm_17_20_L07_B02 | 17 | 20 |
| FVWm_19_20_L07_C02 | 19 | 20 |
| FVWm_17_19_L07_D02 | 17 | 19 |
| FVWm_21_16_L07_E02 | 21 | 16 |
| FVWm_18_16_L07_F02 | 18 | 16 |
| FVWm_18_19_L07_G02 | 18 | 19 |
| FVWm_18_19_L07_H02 | 18 | 19 |
| FVWm_18_17_L07_A03 | 18 | 17 |
| FVWm_18_16_L07_B03 | 18 | 16 |
| FVWm_15_19_L07_C03 | 15 | 19 |
| FVWm_17_18_L07_D03 | 17 | 18 |
| FVWm_18_17_L07_E03 | 18 | 17 |
| FVWm_19_16_L07_F03 | 19 | 16 |
| FVWm_16_17_L07_G03 | 16 | 17 |
| FVWm_15_14_L07_H03 | 15 | 14 |
| FVWm_15_19_L07_A04 | 15 | 19 |
| FVWm_14_19_L07_B04 | 14 | 19 |
| FVWm_22_16_L07_C04 | 22 | 16 |
| FVWm_18_20_L07_F04 | 18 | 20 |
| FVWm_24_15_L07_G04 | 24 | 15 |
| FVWm_19_16_L07_H04 | 19 | 16 |
| FVWm_13_19_L07_B05 | 13 | 19 |
| FVWm_18_19_L07_C05 | 18 | 19 |
| FVWm_16_17_L07_D05 | 16 | 17 |
| FVWm_12_18_L07_E05 | 12 | 18 |
| FVWm_21_21_L07_F05 | 21 | 21 |
| FVWm_19_18_L07_G05 | 19 | 18 |
| FVWm_19_17_L07_H05 | 19 | 17 |
| FVWm_15_15_L07_B06 | 15 | 15 |
| FVWm_19_22_L07_C06 | 19 | 22 |
| FVWm_17_18_L07_D06 | 17 | 18 |
| FVWm_16_15_L07_E06 | 16 | 15 |
| FVWm_17_18_L07_G06 | 17 | 18 |
| FVWm_17_16_L07_A07 | 17 | 16 |
| FVWm_19_17_L07_C07 | 19 | 17 |
| FVWm_13_19_L07_D07 | 13 | 19 |
| FVWm_18_22_L07_E07 | 18 | 22 |
| FVWm_15_19_L07_F07 | 15 | 19 |
| FVWm_22_18_L07_G07 | 22 | 18 |
| FVWm_17_17_L07_H07 | 17 | 17 |
| FVWm_13_17_L07_A08 | 13 | 17 |
| FVWm_18_20_L07_B08 | 18 | 20 |
| FVWm_17_18_L07_C08 | 17 | 18 |
| FVWm_18_22_L07_D08 | 18 | 22 |
| FVWm_19_20_L07_F08 | 19 | 20 |

**S1 Table.** Contd.

| Sample_ID | ABN | SBN | Sample_ID | ABN | SBN |
|---|---|---|---|---|---|
| FVWm_18_17_L02_G10 | 18 | 17 | FVWm_16_17_L07_H08 | 16 | 17 |
| FVWm_17_18_L02_H10 | 17 | 18 | FVWm_19_20_L07_A09 | 19 | 20 |
| FVWm_20_19_L02_A11 | 20 | 19 | FVWm_19_18_L07_B09 | 19 | 18 |
| FVWm_18_20_L02_B11 | 18 | 20 | FVWm_18_17_L07_C09 | 18 | 17 |
| FVWm_18_17_L02_C11 | 18 | 17 | FVWm_17_18_L07_D09 | 17 | 18 |
| FVWm_16_19_L02_D11 | 16 | 19 | FVWm_20_19_L07_E09 | 20 | 19 |
| FVWm_18_17_L02_F11 | 18 | 17 | FVWm_19_20_L07_F09 | 19 | 20 |
| FVWm_22_20_L02_G11 | 22 | 20 | FVWm_16_18_L07_G09 | 16 | 18 |
| FVWm_17_17_L02_H11 | 17 | 17 | FVWm_17_16_L07_H09 | 17 | 16 |
| FVWm_18_16_L02_E12 | 18 | 16 | FVWm_17_18_L07_A10 | 17 | 18 |
| FVWm_19_17_L02_G12 | 19 | 17 | FVWm_16_18_L07_B10 | 16 | 18 |
| FVWm_17_17_L02_H12 | 17 | 17 | FVWm_18_16_L07_D10 | 18 | 16 |
| FVWm_22_22_L04_A01 | 22 | 22 | FVWm_16_20_L07_E10 | 16 | 20 |
| FVWm_18_17_L04_B01 | 18 | 17 | FVWm_20_26_L07_F10 | 20 | 26 |
| FVWm_16_19_L04_C01 | 16 | 19 | FVWm_17_20_L07_G10 | 17 | 20 |
| FVWm_20_19_L04_D01 | 20 | 19 | FVWm_22_21_L07_H10 | 22 | 21 |
| FVWm_18_16_L04_F01 | 18 | 16 | FVWm_17_20_L07_B11 | 17 | 20 |
| FVWm_17_16_L04_G01 | 17 | 16 | FVWm_19_19_L07_C11 | 19 | 19 |
| FVWm_17_19_L04_H01 | 17 | 19 | FVWm_15_18_L07_D11 | 15 | 18 |
| FVWm_19_16_L04_B02 | 19 | 16 | FVWm_21_18_L07_E11 | 21 | 18 |
| FVWm_20_17_L04_C02 | 20 | 17 | FVWm_21_18_L07_F11 | 21 | 18 |
| FVWm_21_19_L04_E02 | 21 | 19 | FVWm_19_16_L07_G11 | 19 | 16 |
| FVWm_16_16_L04_G02 | 16 | 16 | FVWm_15_18_L07_H11 | 15 | 18 |
| FVWm_17_18_L04_H02 | 17 | 18 | FVWm_15_18_L07_A12 | 15 | 18 |
| FVWm_18_16_L04_A03 | 18 | 16 | FVWm_18_18_L07_B12 | 18 | 18 |
| FVWm_19_16_L04_B03 | 19 | 16 | FVWm_18_17_L07_F12 | 18 | 17 |
| FVWm_21_20_L04_C03 | 21 | 20 | FVWm_15_17_L07_G12 | 15 | 17 |
| FVWm_20_18_L04_D03 | 20 | 18 | FVWm_19_15_L08_A01 | 19 | 15 |
| FVWm_18_16_L04_E03 | 18 | 16 | FVWm_19_19_L08_B01 | 19 | 19 |
| FVWm_18_20_L04_F03 | 18 | 20 | FVWm_18_17_L08_C01 | 18 | 17 |
| FVWm_16_18_L04_G03 | 16 | 18 | FVWm_20_20_L08_G01 | 20 | 20 |
| FVWm_17_18_L04_A04 | 17 | 18 | FVWm_17_16_L08_H01 | 17 | 16 |
| FVWm_20_17_L04_B04 | 20 | 17 | FVWm_18_18_L08_C02 | 18 | 18 |
| FVWm_16_17_L04_C04 | 16 | 17 | FVWm_21_18_L08_D02 | 21 | 18 |
| FVWm_17_17_L04_D04 | 17 | 17 | FVWm_18_16_L08_F02 | 18 | 16 |
| FVWm_15_20_L04_G04 | 15 | 20 | FVWm_20_20_L08_G02 | 20 | 20 |
| FVWm_19_17_L04_H04 | 19 | 17 | FVWm_16_18_L08_A03 | 16 | 18 |
| FVWm_15_16_L04_A05 | 15 | 16 | FVWm_18_24_L08_C03 | 18 | 24 |
| FVWm_19_16_L04_B05 | 19 | 16 | FVWm_21_20_L08_F03 | 21 | 20 |
| FVWm_15_17_L04_D05 | 15 | 17 | FVWm_21_17_L08_G03 | 21 | 17 |
| FVWm_15_17_L04_E05 | 15 | 17 | FVWm_15_18_L08_H03 | 15 | 18 |
| FVWm_20_19_L04_F05 | 20 | 19 | FVWm_19_15_L08_A04 | 19 | 15 |
| FVWm_19_20_L04_G05 | 19 | 20 | FVWm_19_17_L08_D04 | 19 | 17 |
| FVWm_18_17_L04_A06 | 18 | 17 | FVWm_19_18_L08_F04 | 19 | 18 |
| FVWm_20_18_L04_C06 | 20 | 18 | FVWm_14_20_L08_B05 | 14 | 20 |
| FVWm_16_19_L04_E06 | 16 | 19 | FVWm_17_14_L08_D05 | 17 | 14 |
| FVWm_22_17_L04_F06 | 22 | 17 | FVWm_18_19_L08_F05 | 18 | 19 |
| FVWm_14_18_L04_G06 | 14 | 18 | FVWm_20_18_L08_G05 | 20 | 18 |
| FVWm_19_18_L04_A07 | 19 | 18 | FVWm_18_17_L08_H05 | 18 | 17 |
| FVWm_16_18_L04_C07 | 16 | 18 | FVWm_15_19_L08_A06 | 15 | 19 |
| FVWm_18_17_L04_D07 | 18 | 17 | FVWm_15_15_L08_B06 | 15 | 15 |
| FVWm_20_16_L04_E07 | 20 | 16 | FVWm_16_18_L08_C06 | 16 | 18 |
| FVWm_21_19_L04_F07 | 21 | 19 | FVWm_18_16_L08_D06 | 18 | 16 |
| FVWm_19_18_L04_H07 | 19 | 18 | FVWm_19_22_L08_E06 | 19 | 22 |
| FVWm_18_18_L04_A08 | 18 | 18 | FVWm_15_18_L08_F06 | 15 | 18 |

**S1 Table.** Contd.

| Sample_ID | ABN | SBN | Sample_ID | ABN | SBN |
|---|---|---|---|---|---|
| FVWm_17_18_L04_D08 | 17 | 18 | FVWm_19_20_L08_G06 | 19 | 20 |
| FVWm_18_18_L04_F08 | 18 | 18 | FVWm_18_23_L08_H06 | 18 | 23 |
| FVWm_16_19_L04_G08 | 16 | 19 | FVWm_18_17_L08_B07 | 18 | 17 |
| FVWm_19_18_L04_H08 | 19 | 18 | FVWm_17_17_L08_C07 | 17 | 17 |
| FVWm_17_18_L04_A09 | 17 | 18 | FVWm_10_19_L08_E07 | 10 | 19 |
| FVWm_19_18_L04_B09 | 19 | 18 | FVWm_17_17_L08_F07 | 17 | 17 |
| FVWm_15_15_L04_C09 | 15 | 15 | FVWm_18_17_L08_H07 | 18 | 17 |
| FVWm_17_18_L04_D09 | 17 | 18 | FVWm_18_18_L08_A08 | 18 | 18 |
| FVWm_17_16_L04_E09 | 17 | 16 | FVWm_15_20_L08_B08 | 15 | 20 |
| FVWm_16_17_L04_F09 | 16 | 17 | FVWm_19_15_L08_C08 | 19 | 15 |
| FVWm_15_18_L04_G09 | 15 | 18 | FVWm_22_14_L08_D08 | 22 | 14 |
| FVWm_17_17_L04_H09 | 17 | 17 | FVWm_17_18_L08_G08 | 17 | 18 |
| FVWm_18_19_L04_A10 | 18 | 19 | FVWm_20_17_L08_A09 | 20 | 17 |
| FVWm_19_17_L04_B10 | 19 | 17 | FVWm_20_19_L08_B09 | 20 | 19 |
| FVWm_20_17_L04_C10 | 20 | 17 | FVWm_18_17_L08_C09 | 18 | 17 |
| FVWm_18_21_L04_D10 | 18 | 21 | FVWm_17_23_L08_F09 | 17 | 23 |
| FVWm_21_20_L04_E10 | 21 | 20 | FVWm_16_20_L08_G09 | 16 | 20 |
| FVWm_17_16_L04_F10 | 17 | 16 | FVWm_17_17_L08_A10 | 17 | 17 |
| FVWm_18_15_L04_G10 | 18 | 15 | FVWm_19_18_L08_B10 | 19 | 18 |
| FVWm_20_20_L04_H10 | 20 | 20 | FVWm_14_19_L08_E10 | 14 | 19 |
| FVWm_20_17_L04_A11 | 20 | 17 | FVWm_20_17_L08_F10 | 20 | 17 |
| FVWm_19_19_L04_B11 | 19 | 19 | FVWm_22_19_L08_G10 | 22 | 19 |
| FVWm_19_18_L04_F11 | 19 | 18 | FVWm_19_16_L08_A11 | 19 | 16 |
| FVWm_16_20_L04_G11 | 16 | 20 | FVWm_08_20_L08_B11 | 08 | 20 |
| FVWm_18_17_L04_B12 | 18 | 17 | FVWm_16_17_L08_C11 | 16 | 17 |
| FVWm_20_16_L04_C12 | 20 | 16 | FVWm_21_18_L08_D11 | 21 | 18 |
| FVWm_21_22_L04_G12 | 21 | 22 | FVWm_20_19_L08_E11 | 20 | 19 |
| FVWm_16_16_L04_H12 | 16 | 16 | FVWm_17_16_L08_F11 | 17 | 16 |
| FVWm_19_18_L12_A01 | 19 | 18 | FVWm_17_17_L08_G11 | 17 | 17 |
| FVWm_17_14_L12_B01 | 17 | 14 | FVWm_18_21_L08_A12 | 18 | 21 |
| FVWm_16_17_L12_C01 | 16 | 17 | FVWm_19_20_L08_F12 | 19 | 20 |
| FVWm_16_19_L12_D01 | 16 | 19 | FVWm_15_19_L08_H12 | 15 | 19 |
| FVWm_14_18_L12_E01 | 14 | 18 | FVWm_17_18_L09_A01 | 17 | 18 |
| FVWm_21_19_L12_F01 | 21 | 19 | FVWm_17_17_L09_B01 | 17 | 17 |
| FVWm_18_19_L12_G01 | 18 | 19 | FVWm_19_18_L09_D01 | 19 | 18 |
| FVWm_15_20_L12_H01 | 15 | 20 | FVWm_18_19_L09_E01 | 18 | 19 |
| FVWm_18_19_L12_A02 | 18 | 19 | FVWm_17_18_L09_F01 | 17 | 18 |
| FVWm_15_22_L12_B02 | 15 | 22 | FVWm_15_16_L09_G01 | 15 | 16 |
| FVWm_17_19_L12_C02 | 17 | 19 | FVWm_18_18_L09_H01 | 18 | 18 |
| FVWm_19_15_L12_D02 | 19 | 15 | FVWm_18_19_L09_A02 | 18 | 19 |
| FVWm_21_21_L12_E02 | 21 | 21 | FVWm_17_18_L09_B02 | 17 | 18 |
| FVWm_16_19_L12_F02 | 16 | 19 | FVWm_17_17_L09_C02 | 17 | 17 |
| FVWm_19_19_L12_G02 | 19 | 19 | FVWm_20_21_L09_D02 | 20 | 21 |
| FVWm_18_15_L12_H02 | 18 | 15 | FVWm_19_17_L09_E02 | 19 | 17 |
| FVWm_17_20_L12_A03 | 17 | 20 | FVWm_18_20_L09_F02 | 18 | 20 |
| FVWm_16_16_L12_B03 | 16 | 16 | FVWm_21_18_L09_G02 | 21 | 18 |
| FVWm_19_20_L12_C03 | 19 | 20 | FVWm_14_19_L09_H02 | 14 | 19 |
| FVWm_19_17_L12_D03 | 19 | 17 | FVWm_20_17_L09_A03 | 20 | 17 |
| FVWm_18_20_L12_E03 | 18 | 20 | FVWm_22_19_L09_B03 | 22 | 19 |
| FVWm_18_19_L12_F03 | 18 | 19 | FVWm_17_20_L09_C03 | 17 | 20 |
| FVWm_16_18_L12_G03 | 16 | 18 | FVWm_16_20_L09_D03 | 16 | 20 |
| FVWm_25_18_L12_H03 | 25 | 18 | FVWm_14_16_L09_E03 | 14 | 16 |
| FVWm_19_18_L12_A04 | 19 | 18 | FVWm_17_21_L09_G03 | 17 | 21 |
| FVWm_14_17_L12_B04 | 14 | 17 | FVWm_18_16_L09_B04 | 18 | 16 |
| FVWm_16_20_L12_C04 | 16 | 20 | FVWm_17_19_L09_C04 | 17 | 19 |

**S1 Table.** Contd.

| Sample_ID | ABN | SBN |
|---|---|---|
| FVWm_17_18_L12_D04 | 17 | 18 |
| FVWm_16_14_L12_E04 | 16 | 14 |
| FVWm_14_16_L12_F04 | 14 | 16 |
| FVWm_18_20_L12_G04 | 18 | 20 |
| FVWm_16_14_L12_H04 | 16 | 14 |
| FVWm_18_19_L12_A05 | 18 | 19 |
| FVWm_16_18_L12_B05 | 16 | 18 |
| FVWm_17_17_L12_C05 | 17 | 17 |
| FVWm_14_21_L12_D05 | 14 | 21 |
| FVWm_20_18_L12_F05 | 20 | 18 |
| FVWm_18_16_L12_G05 | 18 | 16 |
| FVWm_17_16_L12_H05 | 17 | 16 |
| FVWm_16_16_L12_A06 | 16 | 16 |
| FVWm_20_17_L12_B06 | 20 | 17 |
| FVWm_18_20_L12_C06 | 18 | 20 |
| FVWm_17_18_L12_D06 | 17 | 18 |
| FVWm_20_17_L12_E06 | 20 | 17 |
| FVWm_18_18_L12_F06 | 18 | 18 |
| FVWm_21_17_L12_G06 | 21 | 17 |
| FVWm_17_18_L12_H06 | 17 | 18 |
| FVWm_19_18_L12_A07 | 19 | 18 |
| FVWm_15_22_L12_B07 | 15 | 22 |
| FVWm_17_18_L12_C07 | 17 | 18 |
| FVWm_15_19_L12_D07 | 15 | 19 |
| FVWm_17_20_L12_E07 | 17 | 20 |
| FVWm_17_17_L12_F07 | 17 | 17 |
| FVWm_18_19_L12_G07 | 18 | 19 |
| FVWm_18_17_L12_H07 | 18 | 17 |
| FVWm_18_19_L12_A08 | 18 | 19 |
| FVWm_16_17_L12_B08 | 16 | 17 |
| FVWm_21_15_L12_C08 | 21 | 15 |
| FVWm_16_16_L12_D08 | 16 | 16 |
| FVWm_18_18_L12_E08 | 18 | 18 |
| FVWm_19_18_L12_F08 | 19 | 18 |
| FVWm_15_19_L12_G08 | 15 | 19 |
| FVWm_17_17_L12_A09 | 17 | 17 |
| FVWm_20_21_L12_B09 | 20 | 21 |
| FVWm_18_18_L12_C09 | 18 | 18 |
| FVWm_17_18_L12_D09 | 17 | 18 |
| FVWm_20_17_L12_E09 | 20 | 17 |
| FVWm_16_18_L12_F09 | 16 | 18 |
| FVWm_17_17_L12_G09 | 17 | 17 |
| FVWm_15_17_L12_H09 | 15 | 17 |
| FVWm_19_17_L12_A10 | 19 | 17 |
| FVWm_21_19_L12_B10 | 21 | 19 |
| FVWm_20_17_L12_C10 | 20 | 17 |
| FVWm_21_20_L12_D10 | 21 | 20 |
| FVWm_17_17_L12_E10 | 17 | 17 |
| FVWm_19_19_L12_F10 | 19 | 19 |
| FVWm_15_18_L12_G10 | 15 | 18 |
| FVWm_15_19_L12_H10 | 15 | 19 |
| FVWm_19_20_L12_A11 | 19 | 20 |
| FVWm_21_17_L12_B11 | 21 | 17 |
| FVWm_15_19_L12_C11 | 15 | 19 |
| FVWm_18_17_L12_D11 | 18 | 17 |

| Sample_ID | ABN | SBN |
|---|---|---|
| FVWm_19_18_L09_D04 | 19 | 18 |
| FVWm_16_17_L09_E04 | 16 | 17 |
| FVWm_17_21_L09_F04 | 17 | 21 |
| FVWm_20_19_L09_H04 | 20 | 19 |
| FVWm_18_19_L09_B05 | 18 | 19 |
| FVWm_16_17_L09_C05 | 16 | 17 |
| FVWm_18_16_L09_D05 | 18 | 16 |
| FVWm_18_17_L09_E05 | 18 | 17 |
| FVWm_16_16_L09_F05 | 16 | 16 |
| FVWm_16_18_L09_G05 | 16 | 18 |
| FVWm_18_18_L09_H05 | 18 | 18 |
| FVWm_19_20_L09_A06 | 19 | 20 |
| FVWm_18_19_L09_B06 | 18 | 19 |
| FVWm_16_17_L09_C06 | 16 | 17 |
| FVWm_16_17_L09_D06 | 16 | 17 |
| FVWm_19_16_L09_E06 | 19 | 16 |
| FVWm_23_19_L09_F06 | 23 | 19 |
| FVWm_17_17_L09_G06 | 17 | 17 |
| FVWm_18_17_L09_H06 | 18 | 17 |
| FVWm_16_19_L09_A07 | 16 | 19 |
| FVWm_17_17_L09_B07 | 17 | 17 |
| FVWm_18_20_L09_C07 | 18 | 20 |
| FVWm_19_16_L09_D07 | 19 | 16 |
| FVWm_15_16_L09_E07 | 15 | 16 |
| FVWm_18_22_L09_F07 | 18 | 22 |
| FVWm_20_19_L09_G07 | 20 | 19 |
| FVWm_17_18_L09_H07 | 17 | 18 |
| FVWm_15_18_L09_A08 | 15 | 18 |
| FVWm_14_17_L09_B08 | 14 | 17 |
| FVWm_17_23_L09_C08 | 17 | 23 |
| FVWm_19_17_L09_D08 | 19 | 17 |
| FVWm_16_16_L09_E08 | 16 | 16 |
| FVWm_19_17_L09_F08 | 19 | 17 |
| FVWm_17_17_L09_G08 | 17 | 17 |
| FVWm_20_20_L09_H08 | 20 | 20 |
| FVWm_17_16_L09_A09 | 17 | 16 |
| FVWm_18_16_L09_B09 | 18 | 16 |
| FVWm_18_17_L09_C09 | 18 | 17 |
| FVWm_17_19_L09_F09 | 17 | 19 |
| FVWm_17_20_L09_H09 | 17 | 20 |
| FVWm_18_23_L09_A10 | 18 | 23 |
| FVWm_18_17_L09_B10 | 18 | 17 |
| FVWm_18_18_L09_C10 | 18 | 18 |
| FVWm_18_15_L09_D10 | 18 | 15 |
| FVWm_16_21_L09_E10 | 16 | 21 |
| FVWm_18_21_L09_G10 | 18 | 21 |
| FVWm_14_20_L09_H10 | 14 | 20 |
| FVWm_08_18_L09_A11 | 08 | 18 |
| FVWm_20_17_L09_B11 | 20 | 17 |
| FVWm_14_17_L09_C11 | 14 | 17 |
| FVWm_18_18_L09_D11 | 18 | 18 |
| FVWm_14_16_L09_E11 | 14 | 16 |
| FVWm_20_18_L09_F11 | 20 | 18 |
| FVWm_15_20_L09_G11 | 15 | 20 |
| FVWm_19_20_L09_B12 | 19 | 20 |

**S1 Table.** Contd.

| Sample_ID | ABN | SBN | Sample_ID | ABN | SBN |
|---|---|---|---|---|---|
| FVWm_21_18_L12_E11 | 21 | 18 | FVWm_16_15_L09_C12 | 16 | 15 |
| FVWm_22_18_L12_F11 | 22 | 18 | FVWm_18_18_L09_D12 | 18 | 18 |
| FVWm_16_19_L12_G11 | 16 | 19 | FVWm_20_18_L09_E12 | 20 | 18 |
| FVWm_17_17_L12_H11 | 17 | 17 | FVWm_16_16_L09_G12 | 16 | 16 |
| FVWm_20_23_L12_A12 | 20 | 23 | FVWm_19_20_L03_B01 | 19 | 20 |
| FVWm_17_19_L12_B12 | 17 | 19 | FVWm_23_23_L03_C01 | 23 | 23 |
| FVWm_18_18_L12_C12 | 18 | 18 | FVWm_20_18_L03_D01 | 20 | 18 |
| FVWm_16_19_L12_D12 | 16 | 19 | FVWm_16_15_L03_E01 | 16 | 15 |
| FVWm_17_18_L12_E12 | 17 | 18 | FVWm_16_15_L03_G01 | 16 | 15 |
| FVWm_18_17_L12_F12 | 18 | 17 | FVWm_22_18_L03_H01 | 22 | 18 |
| FVWm_21_17_L12_G12 | 21 | 17 | FVWm_19_17_L03_C02 | 19 | 17 |
| FVWm_13_18_L12_H12 | 13 | 18 | FVWm_18_20_L03_D02 | 18 | 20 |
| FVWm_17_22_L01_A01 | 17 | 22 | FVWm_17_16_L03_E02 | 17 | 16 |
| FVWm_19_18_L01_B01 | 19 | 18 | FVWm_19_16_L03_F02 | 19 | 16 |
| FVWm_19_19_L01_C01 | 19 | 19 | FVWm_21_20_L03_G02 | 21 | 20 |
| FVWm_19_16_L01_D01 | 19 | 16 | FVWm_21_19_L03_H02 | 21 | 19 |
| FVWm_20_18_L01_E01 | 20 | 18 | FVWm_17_20_L03_A03 | 17 | 20 |
| FVWm_16_19_L01_F01 | 16 | 19 | FVWm_15_19_L03_B03 | 15 | 19 |
| FVWm_17_19_L01_G01 | 17 | 19 | FVWm_16_19_L03_C03 | 16 | 19 |
| FVWm_19_17_L01_H01 | 19 | 17 | FVWm_18_20_L03_D03 | 18 | 20 |
| FVWm_19_23_L01_A02 | 19 | 23 | FVWm_17_20_L03_E03 | 17 | 20 |
| FVWm_22_21_L01_C02 | 22 | 21 | FVWm_16_16_L03_F03 | 16 | 16 |
| FVWm_21_17_L01_E02 | 21 | 17 | FVWm_20_18_L03_G03 | 20 | 18 |
| FVWm_20_22_L01_F02 | 20 | 22 | FVWm_14_16_L03_A04 | 14 | 16 |
| FVWm_16_17_L01_G02 | 16 | 17 | FVWm_21_20_L03_D04 | 21 | 20 |
| FVWm_21_18_L01_A03 | 21 | 18 | FVWm_18_21_L03_E04 | 18 | 21 |
| FVWm_20_18_L01_B03 | 20 | 18 | FVWm_17_17_L03_G04 | 17 | 17 |
| FVWm_18_20_L01_C03 | 18 | 20 | FVWm_18_19_L03_H04 | 18 | 19 |
| FVWm_14_19_L01_D03 | 14 | 19 | FVWm_16_19_L03_A05 | 16 | 19 |
| FVWm_14_18_L01_E03 | 14 | 18 | FVWm_19_17_L03_B05 | 19 | 17 |
| FVWm_15_18_L01_F03 | 15 | 18 | FVWm_19_18_L03_D05 | 19 | 18 |
| FVWm_16_20_L01_G03 | 16 | 20 | FVWm_20_17_L03_E05 | 20 | 17 |
| FVWm_16_18_L01_A04 | 16 | 18 | FVWm_17_20_L03_F05 | 17 | 20 |
| FVWm_17_16_L01_B04 | 17 | 16 | FVWm_18_16_L03_G05 | 18 | 16 |
| FVWm_18_18_L01_C04 | 18 | 18 | FVWm_20_18_L03_H05 | 20 | 18 |
| FVWm_19_19_L01_D04 | 19 | 19 | FVWm_20_20_L03_A06 | 20 | 20 |
| FVWm_19_17_L01_F04 | 19 | 17 | FVWm_18_18_L03_B06 | 18 | 18 |
| FVWm_18_18_L01_H04 | 18 | 18 | FVWm_20_18_L03_C06 | 20 | 18 |
| FVWm_19_16_L01_A05 | 19 | 16 | FVWm_17_17_L03_E06 | 17 | 17 |
| FVWm_17_17_L01_B05 | 17 | 17 | FVWm_18_19_L03_F06 | 18 | 19 |
| FVWm_17_16_L01_D05 | 17 | 16 | FVWm_17_21_L03_G06 | 17 | 21 |
| FVWm_17_16_L01_F05 | 17 | 16 | FVWm_16_18_L03_H06 | 16 | 18 |
| FVWm_21_15_L01_G05 | 21 | 15 | FVWm_16_17_L03_A07 | 16 | 17 |
| FVWm_18_16_L01_H05 | 18 | 16 | FVWm_18_19_L03_B07 | 18 | 19 |
| FVWm_20_20_L01_A06 | 20 | 20 | FVWm_15_25_L03_D07 | 15 | 25 |
| FVWm_17_18_L01_B06 | 17 | 18 | FVWm_17_16_L03_E07 | 17 | 16 |
| FVWm_19_23_L01_C06 | 19 | 23 | FVWm_17_19_L03_F07 | 17 | 19 |
| FVWm_17_19_L01_E06 | 17 | 19 | FVWm_19_17_L03_G07 | 19 | 17 |
| FVWm_20_18_L01_F06 | 20 | 18 | FVWm_17_18_L03_H07 | 17 | 18 |
| FVWm_18_21_L01_G06 | 18 | 21 | FVWm_21_17_L03_A08 | 21 | 17 |
| FVWm_18_20_L01_H06 | 18 | 20 | FVWm_18_17_L03_D08 | 18 | 17 |
| FVWm_16_16_L01_A07 | 16 | 16 | FVWm_16_21_L03_E08 | 16 | 21 |
| FVWm_19_18_L01_B07 | 19 | 18 | FVWm_18_20_L03_F08 | 18 | 20 |
| FVWm_16_17_L01_C07 | 16 | 17 | FVWm_19_18_L03_G08 | 19 | 18 |
| FVWm_16_16_L01_D07 | 16 | 16 | FVWm_17_19_L03_H08 | 17 | 19 |

**S1 Table.** Contd.

| Sample_ID | ABN | SBN |
|---|---|---|
| FVWm_17_19_L01_E07 | 17 | 19 |
| FVWm_19_18_L01_F07 | 19 | 18 |
| FVWm_21_19_L01_G07 | 21 | 19 |
| FVWm_19_20_L01_H07 | 19 | 20 |
| FVWm_15_18_L01_A08 | 15 | 18 |
| FVWm_18_17_L01_B08 | 18 | 17 |
| FVWm_22_20_L01_C08 | 22 | 20 |
| FVWm_18_17_L01_D08 | 18 | 17 |
| FVWm_17_19_L01_F08 | 17 | 19 |
| FVWm_20_19_L01_G08 | 20 | 19 |
| FVWm_20_20_L01_H08 | 20 | 20 |
| FVWm_21_19_L01_B09 | 21 | 19 |
| FVWm_18_18_L01_E09 | 18 | 18 |
| FVWm_18_21_L01_F09 | 18 | 21 |
| FVWm_17_21_L01_G09 | 17 | 21 |
| FVWm_21_20_L01_H09 | 21 | 20 |
| FVWm_19_17_L01_A10 | 19 | 17 |
| FVWm_18_17_L01_C10 | 18 | 17 |
| FVWm_16_18_L01_D10 | 16 | 18 |
| FVWm_21_21_L01_E10 | 21 | 21 |
| FVWm_17_18_L01_F10 | 17 | 18 |
| FVWm_20_20_L01_H10 | 20 | 20 |
| FVWm_21_21_L01_A11 | 21 | 21 |
| FVWm_20_17_L01_B11 | 20 | 17 |
| FVWm_17_15_L01_C11 | 17 | 15 |
| FVWm_17_18_L01_D11 | 17 | 18 |
| FVWm_17_19_L01_E11 | 17 | 19 |
| FVWm_19_17_L01_F11 | 19 | 17 |
| FVWm_20_20_L01_G11 | 20 | 20 |
| FVWm_19_17_L01_H11 | 19 | 17 |
| FVWm_19_19_L01_B12 | 19 | 19 |
| FVWm_18_17_L01_E12 | 18 | 17 |
| FVWm_17_19_L01_F12 | 17 | 19 |
| FVWm_17_17_L01_G12 | 17 | 17 |
| FVWm_22_19_L01_H12 | 22 | 19 |
| FVWm_20_18_L05_A01 | 20 | 18 |
| FVWm_20_18_L05_B01 | 20 | 18 |
| FVWm_21_17_L05_C01 | 21 | 17 |
| FVWm_14_19_L05_F01 | 14 | 19 |
| FVWm_17_21_L05_G01 | 17 | 21 |
| FVWm_21_17_L05_H01 | 21 | 17 |
| FVWm_16_20_L05_D02 | 16 | 20 |
| FVWm_15_16_L05_E02 | 15 | 16 |
| FVWm_19_20_L05_F02 | 19 | 20 |
| FVWm_18_20_L05_G02 | 18 | 20 |
| FVWm_22_20_L05_H02 | 22 | 20 |
| FVWm_19_17_L05_B03 | 19 | 17 |
| FVWm_21_16_L05_C03 | 21 | 16 |
| FVWm_20_15_L05_E03 | 20 | 15 |
| FVWm_17_18_L05_G03 | 17 | 18 |
| FVWm_17_20_L05_H03 | 17 | 20 |
| FVWm_18_18_L05_A04 | 18 | 18 |
| FVWm_17_14_L05_B04 | 17 | 14 |
| FVWm_18_20_L05_D04 | 18 | 20 |
| FVWm_18_16_L05_G04 | 18 | 16 |

| Sample_ID | ABN | SBN |
|---|---|---|
| FVWm_19_19_L03_A09 | 19 | 19 |
| FVWm_18_18_L03_B09 | 18 | 18 |
| FVWm_20_19_L03_D09 | 20 | 19 |
| FVWm_18_19_L03_E09 | 18 | 19 |
| FVWm_19_18_L03_F09 | 19 | 18 |
| FVWm_18_18_L03_G09 | 18 | 18 |
| FVWm_19_16_L03_H09 | 19 | 16 |
| FVWm_15_18_L03_A10 | 15 | 18 |
| FVWm_19_18_L03_F10 | 19 | 18 |
| FVWm_13_17_L03_G10 | 13 | 17 |
| FVWm_18_16_L03_H10 | 18 | 16 |
| FVWm_17_22_L03_A11 | 17 | 22 |
| FVWm_18_17_L03_C11 | 18 | 17 |
| FVWm_16_16_L03_D11 | 16 | 16 |
| FVWm_17_20_L03_A12 | 17 | 20 |
| FVWm_19_19_L03_B12 | 19 | 19 |
| FVWm_16_16_L03_C12 | 16 | 16 |
| FVWm_21_17_L03_D12 | 21 | 17 |
| FVWm_18_18_L03_F12 | 18 | 18 |
| FVWm_22_16_L03_G12 | 22 | 16 |
| FVWm_17_19_L03_H12 | 17 | 19 |
| FVWm_18_20_L10_A01 | 18 | 20 |
| FVWm_17_20_L10_B01 | 17 | 20 |
| FVWm_16_20_L10_C01 | 16 | 20 |
| FVWm_18_22_L10_D01 | 18 | 22 |
| FVWm_16_17_L10_E01 | 16 | 17 |
| FVWm_13_14_L10_F01 | 13 | 14 |
| FVWm_18_19_L10_A02 | 18 | 19 |
| FVWm_17_19_L10_B02 | 17 | 19 |
| FVWm_19_19_L10_C02 | 19 | 19 |
| FVWm_18_18_L10_D02 | 18 | 18 |
| FVWm_14_19_L10_E02 | 14 | 19 |
| FVWm_17_20_L10_F02 | 17 | 20 |
| FVWm_19_20_L10_G02 | 19 | 20 |
| FVWm_17_20_L10_H02 | 17 | 20 |
| FVWm_16_18_L10_A03 | 16 | 18 |
| FVWm_17_23_L10_B03 | 17 | 23 |
| FVWm_17_18_L10_C03 | 17 | 18 |
| FVWm_16_16_L10_D03 | 16 | 16 |
| FVWm_19_18_L10_E03 | 19 | 18 |
| FVWm_18_20_L10_F03 | 18 | 20 |
| FVWm_17_18_L10_G03 | 17 | 18 |
| FVWm_17_16_L10_H03 | 17 | 16 |
| FVWm_16_19_L10_A04 | 16 | 19 |
| FVWm_18_17_L10_B04 | 18 | 17 |
| FVWm_15_17_L10_C04 | 15 | 17 |
| FVWm_15_17_L10_D04 | 15 | 17 |
| FVWm_19_20_L10_E04 | 19 | 20 |
| FVWm_15_18_L10_F04 | 15 | 18 |
| FVWm_18_14_L10_G04 | 18 | 14 |
| FVWm_14_18_L10_H04 | 14 | 18 |
| FVWm_16_18_L10_A05 | 16 | 18 |
| FVWm_16_18_L10_B05 | 16 | 18 |
| FVWm_21_18_L10_C05 | 21 | 18 |
| FVWm_19_17_L10_D05 | 19 | 17 |

**S1 Table.** Contd.

| Sample_ID | ABN | SBN |
|---|---|---|
| FVWm_17_17_L05_B05 | 17 | 17 |
| FVWm_15_20_L05_C05 | 15 | 20 |
| FVWm_18_18_L05_D05 | 18 | 18 |
| FVWm_16_17_L05_E05 | 16 | 17 |
| FVWm_18_20_L05_F05 | 18 | 20 |
| FVWm_14_18_L05_G05 | 14 | 18 |
| FVWm_18_17_L05_H05 | 18 | 17 |
| FVWm_15_16_L05_A06 | 15 | 16 |
| FVWm_18_21_L05_B06 | 18 | 21 |
| FVWm_14_17_L05_C06 | 14 | 17 |
| FVWm_17_18_L05_F06 | 17 | 18 |
| FVWm_18_16_L05_G06 | 18 | 16 |
| FVWm_18_17_L05_H06 | 18 | 17 |
| FVWm_18_16_L05_A07 | 18 | 16 |
| FVWm_20_16_L05_B07 | 20 | 16 |
| FVWm_19_19_L05_C07 | 19 | 19 |
| FVWm_16_18_L05_D07 | 16 | 18 |
| FVWm_17_16_L05_E07 | 17 | 16 |
| FVWm_18_19_L05_F07 | 18 | 19 |
| FVWm_19_20_L05_H07 | 19 | 20 |
| FVWm_14_16_L05_B08 | 14 | 16 |
| FVWm_20_21_L05_C08 | 20 | 21 |
| FVWm_21_16_L05_D08 | 21 | 16 |
| FVWm_13_21_L05_E08 | 13 | 21 |
| FVWm_21_16_L05_F08 | 21 | 16 |
| FVWm_17_19_L05_H08 | 17 | 19 |
| FVWm_19_20_L05_A09 | 19 | 20 |
| FVWm_19_17_L05_C09 | 19 | 17 |
| FVWm_16_18_L05_D09 | 16 | 18 |
| FVWm_15_21_L05_E09 | 15 | 21 |
| FVWm_17_19_L05_F09 | 17 | 19 |
| FVWm_20_19_L05_G09 | 20 | 19 |
| FVWm_22_20_L05_H09 | 22 | 20 |
| FVWm_18_18_L05_A10 | 18 | 18 |
| FVWm_18_19_L05_D10 | 18 | 19 |
| FVWm_22_20_L05_F10 | 22 | 20 |
| FVWm_20_16_L05_G10 | 20 | 16 |
| FVWm_16_16_L05_H10 | 16 | 16 |
| FVWm_16_15_L05_A11 | 16 | 15 |
| FVWm_21_15_L05_B11 | 21 | 15 |
| FVWm_17_21_L05_C11 | 17 | 21 |
| FVWm_16_20_L05_D11 | 16 | 20 |
| FVWm_18_18_L05_E11 | 18 | 18 |
| FVWm_17_18_L05_F11 | 17 | 18 |
| FVWm_17_18_L05_G11 | 17 | 18 |
| FVWm_17_17_L05_A12 | 17 | 17 |
| FVWm_19_19_L05_C12 | 19 | 19 |
| FVWm_18_18_L05_D12 | 18 | 18 |
| FVWm_17_19_L05_E12 | 17 | 19 |
| FVWm_16_17_L05_F12 | 16 | 17 |
| FVWm_17_21_L05_G12 | 17 | 21 |
| FVWm_19_16_L06_A01 | 19 | 16 |
| FVWm_17_16_L06_B01 | 17 | 16 |
| FVWm_17_17_L06_C01 | 17 | 17 |
| FVWm_17_19_L06_D01 | 17 | 19 |

| Sample_ID | ABN | SBN |
|---|---|---|
| FVWm_22_24_L10_E05 | 22 | 24 |
| FVWm_18_20_L10_F05 | 18 | 20 |
| FVWm_16_19_L10_G05 | 16 | 19 |
| FVWm_21_19_L10_H05 | 21 | 19 |
| FVWm_20_19_L10_A06 | 20 | 19 |
| FVWm_20_19_L10_B06 | 20 | 19 |
| FVWm_16_18_L10_C06 | 16 | 18 |
| FVWm_21_18_L10_D06 | 21 | 18 |
| FVWm_17_23_L10_E06 | 17 | 23 |
| FVWm_18_19_L10_F06 | 18 | 19 |
| FVWm_17_20_L10_G06 | 17 | 20 |
| FVWm_17_19_L10_H06 | 17 | 19 |
| FVWm_20_18_L10_A07 | 20 | 18 |
| FVWm_14_17_L10_B07 | 14 | 17 |
| FVWm_19_16_L10_C07 | 19 | 16 |
| FVWm_19_16_L10_D07 | 19 | 16 |
| FVWm_17_19_L10_E07 | 17 | 19 |
| FVWm_22_22_L10_F07 | 22 | 22 |
| FVWm_19_21_L10_G07 | 19 | 21 |
| FVWm_20_17_L10_A08 | 20 | 17 |
| FVWm_16_15_L10_B08 | 16 | 15 |
| FVWm_17_19_L10_C08 | 17 | 19 |
| FVWm_17_18_L10_D08 | 17 | 18 |
| FVWm_18_18_L10_E08 | 18 | 18 |
| FVWm_16_18_L10_F08 | 16 | 18 |
| FVWm_18_18_L10_G08 | 18 | 18 |
| FVWm_20_17_L10_H08 | 20 | 17 |
| FVWm_14_16_L10_A09 | 14 | 16 |
| FVWm_18_17_L10_B09 | 18 | 17 |
| FVWm_21_20_L10_C09 | 21 | 20 |
| FVWm_18_19_L10_D09 | 18 | 19 |
| FVWm_19_18_L10_E09 | 19 | 18 |
| FVWm_16_19_L10_F09 | 16 | 19 |
| FVWm_20_17_L10_G09 | 20 | 17 |
| FVWm_15_18_L10_H09 | 15 | 18 |
| FVWm_19_19_L10_A10 | 19 | 19 |
| FVWm_16_18_L10_B10 | 16 | 18 |
| FVWm_23_18_L10_C10 | 23 | 18 |
| FVWm_13_18_L10_D10 | 13 | 18 |
| FVWm_15_19_L10_E10 | 15 | 19 |
| FVWm_22_18_L10_F10 | 22 | 18 |
| FVWm_20_19_L10_G10 | 20 | 19 |
| FVWm_14_18_L10_H10 | 14 | 18 |
| FVWm_17_19_L10_A11 | 17 | 19 |
| FVWm_14_17_L10_B11 | 14 | 17 |
| FVWm_15_17_L10_C11 | 15 | 17 |
| FVWm_17_19_L10_E11 | 17 | 19 |
| FVWm_18_20_L10_F11 | 18 | 20 |
| FVWm_18_18_L10_G11 | 18 | 18 |
| FVWm_19_20_L10_H11 | 19 | 20 |
| FVWm_17_17_L10_A12 | 17 | 17 |
| FVWm_19_17_L10_C12 | 19 | 17 |
| FVWm_19_16_L10_E12 | 19 | 16 |
| FVWm_19_18_L10_F12 | 19 | 18 |
| FVWm_17_18_L10_G12 | 17 | 18 |

**S1 Table.** Contd.

| Sample_ID | ABN | SBN | Sample_ID | ABN | SBN |
|---|---|---|---|---|---|
| FVWm_19_19_L06_F01 | 19 | 19 | FVWm_18_16_L11_A01 | 18 | 16 |
| FVWm_18_16_L06_G01 | 18 | 16 | FVWm_17_19_L11_B01 | 17 | 19 |
| FVWm_19_20_L06_H01 | 19 | 20 | FVWm_17_18_L11_C01 | 17 | 18 |
| FVWm_15_19_L06_A02 | 15 | 19 | FVWm_16_20_L11_F01 | 16 | 20 |
| FVWm_14_18_L06_B02 | 14 | 18 | FVWm_19_20_L11_G01 | 19 | 20 |
| FVWm_21_17_L06_C02 | 21 | 17 | FVWm_19_15_L11_H01 | 19 | 15 |
| FVWm_19_18_L06_D02 | 19 | 18 | FVWm_21_17_L11_C02 | 21 | 17 |
| FVWm_19_16_L06_F02 | 19 | 16 | FVWm_20_20_L11_G02 | 20 | 20 |
| FVWm_17_18_L06_G02 | 17 | 18 | FVWm_18_16_L11_H02 | 18 | 16 |
| FVWm_16_17_L06_A03 | 16 | 17 | FVWm_17_15_L11_A03 | 17 | 15 |
| FVWm_17_16_L06_C03 | 17 | 16 | FVWm_20_16_L11_B03 | 20 | 16 |
| FVWm_18_20_L06_E03 | 18 | 20 | FVWm_20_19_L11_C03 | 20 | 19 |
| FVWm_20_17_L06_F03 | 20 | 17 | FVWm_15_18_L11_D03 | 15 | 18 |
| FVWm_20_18_L06_G03 | 20 | 18 | FVWm_20_18_L11_E03 | 20 | 18 |
| FVWm_18_20_L06_H03 | 18 | 20 | FVWm_19_20_L11_F03 | 19 | 20 |
| FVWm_18_18_L06_A04 | 18 | 18 | FVWm_18_21_L11_G03 | 18 | 21 |
| FVWm_15_20_L06_B04 | 15 | 20 | FVWm_18_22_L11_H03 | 18 | 22 |
| FVWm_20_18_L06_C04 | 20 | 18 | FVWm_17_19_L11_A04 | 17 | 19 |
| FVWm_20_18_L06_D04 | 20 | 18 | FVWm_17_19_L11_B04 | 17 | 19 |
| FVWm_18_18_L06_E04 | 18 | 18 | FVWm_18_17_L11_C04 | 18 | 17 |
| FVWm_18_17_L06_F04 | 18 | 17 | FVWm_17_17_L11_D04 | 17 | 17 |
| FVWm_17_20_L06_G04 | 17 | 20 | FVWm_19_19_L11_E04 | 19 | 19 |
| FVWm_19_16_L06_H04 | 19 | 16 | FVWm_17_17_L11_F04 | 17 | 17 |
| FVWm_21_17_L06_C05 | 21 | 17 | FVWm_18_17_L11_G04 | 18 | 17 |
| FVWm_17_17_L06_D05 | 17 | 17 | FVWm_16_19_L11_H04 | 16 | 19 |
| FVWm_18_20_L06_E05 | 18 | 20 | FVWm_18_15_L11_B05 | 18 | 15 |
| FVWm_16_13_L06_H05 | 16 | 13 | FVWm_17_18_L11_C05 | 17 | 18 |
| FVWm_22_18_L06_A06 | 22 | 18 | FVWm_16_17_L11_D05 | 16 | 17 |
| FVWm_16_15_L06_B06 | 16 | 15 | FVWm_19_18_L11_E05 | 19 | 18 |
| FVWm_17_18_L06_C06 | 17 | 18 | FVWm_17_18_L11_F05 | 17 | 18 |
| FVWm_18_17_L06_D06 | 18 | 17 | FVWm_18_18_L11_G05 | 18 | 18 |
| FVWm_17_17_L06_E06 | 17 | 17 | FVWm_17_21_L11_H05 | 17 | 21 |
| FVWm_19_19_L06_G06 | 19 | 19 | FVWm_16_18_L11_A06 | 16 | 18 |
| FVWm_15_13_L06_H06 | 15 | 13 | FVWm_15_15_L11_B06 | 15 | 15 |
| FVWm_18_17_L06_A07 | 18 | 17 | FVWm_16_16_L11_C06 | 16 | 16 |
| FVWm_22_15_L06_D07 | 22 | 15 | FVWm_19_16_L11_D06 | 19 | 16 |
| FVWm_17_14_L06_E07 | 17 | 14 | FVWm_16_17_L11_E06 | 16 | 17 |
| FVWm_17_18_L06_F07 | 17 | 18 | FVWm_20_20_L11_F06 | 20 | 20 |
| FVWm_17_16_L06_H07 | 17 | 16 | FVWm_18_15_L11_G06 | 18 | 15 |
| FVWm_20_17_L06_A08 | 20 | 17 | FVWm_20_20_L11_H06 | 20 | 20 |
| FVWm_20_17_L06_C08 | 20 | 17 | FVWm_17_18_L11_A07 | 17 | 18 |
| FVWm_19_18_L06_D08 | 19 | 18 | FVWm_16_20_L11_B07 | 16 | 20 |
| FVWm_17_17_L06_E08 | 17 | 17 | FVWm_16_20_L11_C07 | 16 | 20 |
| FVWm_19_19_L06_F08 | 19 | 19 | FVWm_17_17_L11_D07 | 17 | 17 |
| FVWm_19_17_L06_G08 | 19 | 17 | FVWm_18_18_L11_E07 | 18 | 18 |
| FVWm_17_19_L06_A09 | 17 | 19 | FVWm_17_20_L11_F07 | 17 | 20 |
| FVWm_18_18_L06_B09 | 18 | 18 | FVWm_17_18_L11_H07 | 17 | 18 |
| FVWm_20_17_L06_E09 | 20 | 17 | FVWm_21_21_L11_A08 | 21 | 21 |
| FVWm_23_19_L06_F09 | 23 | 19 | FVWm_18_18_L11_B08 | 18 | 18 |
| FVWm_16_17_L06_G09 | 16 | 17 | FVWm_20_17_L11_C08 | 20 | 17 |
| FVWm_18_21_L06_H09 | 18 | 21 | FVWm_19_18_L11_D08 | 19 | 18 |
| FVWm_18_14_L06_A10 | 18 | 14 | FVWm_17_19_L11_E08 | 17 | 19 |
| FVWm_17_17_L06_B10 | 17 | 17 | FVWm_18_15_L11_F08 | 18 | 15 |
| FVWm_20_20_L06_F10 | 20 | 20 | FVWm_19_18_L11_G08 | 19 | 18 |
| FVWm_17_18_L06_H10 | 17 | 18 | FVWm_18_24_L11_H08 | 18 | 24 |

**S1 Table.** Contd.

| Sample_ID | ABN | SBN |
| --- | --- | --- |
| FVWm_19_21_L06_A11 | 19 | 21 |
| FVWm_16_18_L06_B11 | 16 | 18 |
| FVWm_19_21_L06_D11 | 19 | 21 |
| FVWm_19_19_L06_E11 | 19 | 19 |
| FVWm_20_17_L06_F11 | 20 | 17 |
| FVWm_19_20_L06_H11 | 19 | 20 |
| FVWm_18_17_L06_A12 | 18 | 17 |
| FVWm_16_20_L06_B12 | 16 | 20 |
| FVWm_19_15_L06_C12 | 19 | 15 |
| FVWm_18_18_L06_F12 | 18 | 18 |

| Sample_ID | ABN | SBN |
| --- | --- | --- |
| FVWm_18_17_L11_A09 | 18 | 17 |
| FVWm_18_19_L11_B09 | 18 | 19 |
| FVWm_18_21_L11_C09 | 18 | 21 |
| FVWm_19_16_L11_E09 | 19 | 16 |
| FVWm_16_16_L11_F09 | 16 | 16 |
| FVWm_17_20_L11_H09 | 17 | 20 |
| FVWm_15_19_L11_A10 | 15 | 19 |
| FVWm_16_19_L11_B10 | 16 | 19 |
| FVWm_18_18_L11_C10 | 18 | 18 |
