## Supplementary material for "A genomewide association study for bristle number variation in *Drosophila melanogaster*": S1_Text.pdf

**S1 Text.** Ingredients and brief protocol for Macdonald lab cornmeal-yeast-molasses fly media.

28.5-liters water
318-g agar (Genesee Scientific; 66-111. Amount is based on a gel strength of 960 g/cm2, and will change depending on the agar batch)

- Add water to steam kettle, turn on electric mixer, and slowly add agar
- Bring mix to a boil

3,200-ml molasses (Genesee Scientific; 62-117)

- Reduce the kettle pressure to reduce the heat slightly
- Add molasses, and bring mix back to a boil

4-liters water
1,460-g inactive dry yeast (Genesee Scientific; 62-107)

- Mix in bucket using paint-stirring drill attachment

4-liters water
2,600-g yellow cornmeal (Genesee Scientific; 62-101)

- Mix in bucket using paint-stirring drill attachment
- Add both the water/yeast and water/cornmeal mixes to the steam kettle
- Bring mix back to boil, and simmer for ~15-min
- Release pressure from steam kettle, but continue to stir with electric mixer

330-ml water
259-ml propionic acid (ThermoFisher; A258-500)
31-ml phosphoric acid (85%; ThermoFisher; A242-500)

- Pour mix into kettle

400-ml 95% ethanol
1.5-g tegosept (Genesee Scientific; 20-258)

- Dissolve tegosept in ethanol
- Pour mix into kettle
- Fill vials/bottles
