## Supplementary material for "A genomewide association study for bristle number variation in *Drosophila melanogaster*": S2_Fig.pdf

**S2 Fig.** Linkage Disequilibrium (LD) at each cluster of GWAS hits. Plots of LD (in $R^2$) between the lead variant (that with the lowest *P*-value) at each location and all other variants within 3-Mb of it.

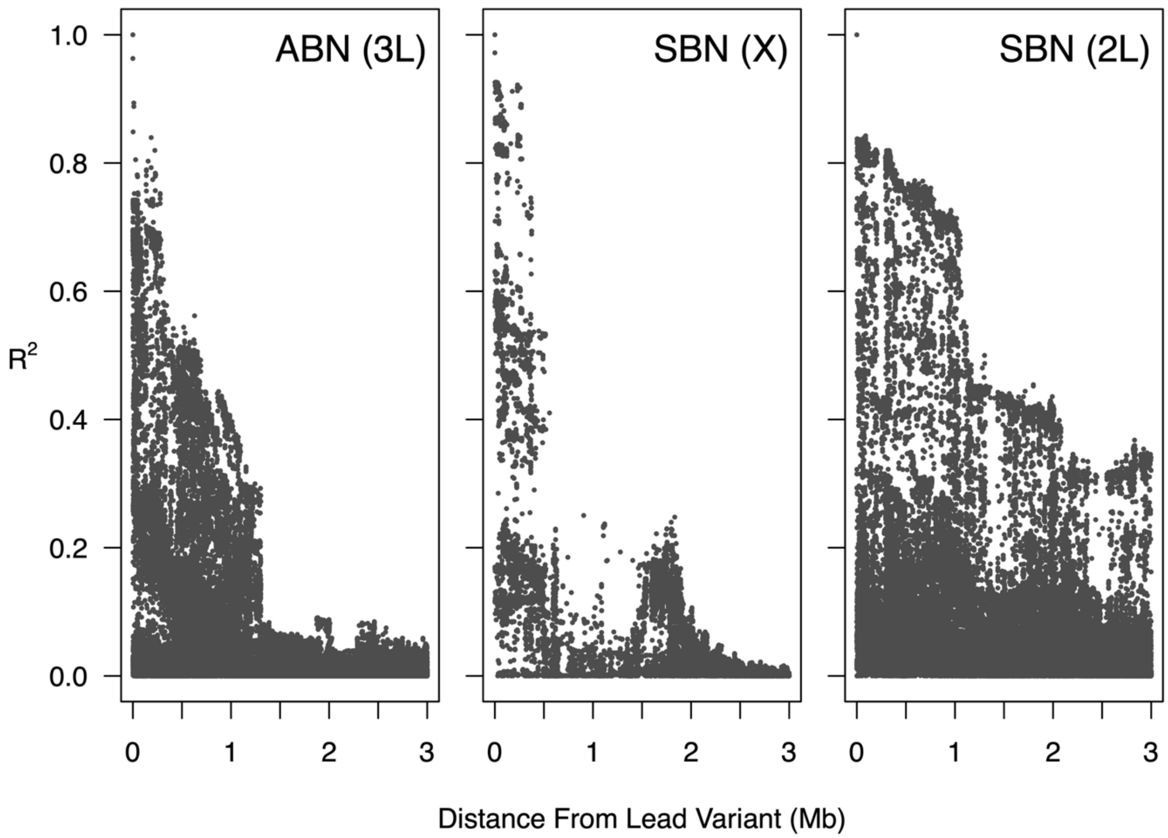
