## Supplementary material for "A genomewide association study for bristle number variation in *Drosophila melanogaster*": S2_Table.pdf

**S2 Table.** Unique dual indexing oligos used for sequencing library amplification and barcoding. The 8-nt index sequences below fit within the longer PCR oligos as follows:

i5_Sequence: 5' - AATGATACGGCGACCACCGAGATCTACACnnnnnnnnTCGTCGGCAGCGTC - 3'
i7_Sequence: 5' - CAAGCAGAAGACGGCATACGAGATnnnnnnnnGTCTCGTGGGCTCGG - 3'

| Plate | Well | Index_ Pair_ID | i5_Sequence | i7_Sequence |
|---|---|---|---|---|
| 1 | A01 | 001 | ATATGCGC | ACGATCAG |
| 1 | A02 | 002 | TGGTACAG | TCGAGAGT |
| 1 | A03 | 003 | AACCGTTC | CTAGCTCA |
| 1 | A04 | 004 | TAACCGGT | ATCGTCTC |
| 1 | A05 | 005 | GAACATCG | TCGACAAG |
| 1 | A06 | 006 | CCTTGTAG | CCTTGGAA |
| 1 | A07 | 007 | TCAGGCTT | ATCATGCG |
| 1 | A08 | 008 | GTTCTCGT | TGTTCCGT |
| 1 | A09 | 009 | AGAACGAG | ATTAGCCG |
| 1 | A10 | 010 | TGCTTCCA | CGATCGAT |
| 1 | A11 | 011 | CTTCGACT | GATCTTGC |
| 1 | A12 | 012 | CACCTGTT | AGGATAGC |
| 1 | B01 | 013 | ATCACACG | GTAGCGTA |
| 1 | B02 | 014 | CCGTAAGA | AGAGTCCA |
| 1 | B03 | 015 | TACGCCTT | GCTACTCT |
| 1 | B04 | 016 | CGACGTTA | CTCTGGAT |
| 1 | B05 | 017 | ATGCACGA | AGATCGTC |
| 1 | B06 | 018 | CCTGATTG | GCTCAGTT |
| 1 | B07 | 019 | GTAGGAGT | GTCCTAAG |
| 1 | B08 | 020 | ACTAGGAG | TATGGCAC |
| 1 | B09 | 021 | CACTAGCT | TCGGATTC |
| 1 | B10 | 022 | ACGACTTG | AACAGCGA |
| 1 | B11 | 023 | CGTGTGTA | CCAACGAA |
| 1 | B12 | 024 | GTTGACCT | CAGTGCTT |
| 1 | C01 | 025 | ACTCCATC | GATCAAGG |
| 1 | C02 | 026 | CAATGTGG | TCTTCGAC |
| 1 | C03 | 027 | TTGCAGAC | ATCGTGGT |
| 1 | C04 | 028 | CAGTCCAA | CGGTAATC |
| 1 | C05 | 029 | ACGTTCAG | AGTTGTGC |
| 1 | C06 | 030 | AACGTCTG | AATGACGC |
| 1 | C07 | 031 | TATCGGTC | TACCGGAT |
| 1 | C08 | 032 | CGCTCTAT | TTGCAACG |
| 1 | C09 | 033 | GATTGCTC | CACTTCAC |
| 1 | C10 | 034 | GATGTGTG | TAGCCATG |
| 1 | C11 | 035 | CGCAATCT | ACAGGCAT |
| 1 | C12 | 036 | TGGTAGCT | AGGTGTTG |
| 1 | D01 | 037 | GATAGGCT | CAGTCACA |
| 1 | D02 | 038 | AGTGGATC | TCGATGAC |
| 1 | D03 | 039 | TTGGACGT | GAAGTGCT |
| 1 | D04 | 040 | ATGACGTC | CTTCCTTC |
| 1 | D05 | 041 | GAAGTTGG | CGAACAAC |
| 1 | D06 | 042 | CATACCAC | AACAACCG |
| 1 | D07 | 043 | CTGTTGAC | ACCTCAGT |
| 1 | D08 | 044 | TGGCATGT | CGTCTTCA |
| 1 | D09 | 045 | ATCGCCAT | TGCGTAAC |
| 1 | D10 | 046 | TTGCGAAG | AACACGCT |
| 1 | D11 | 047 | AGTTCGTC | ACTCGATC |
| 1 | D12 | 048 | GAGCAGTA | TGAGCTGT |
| 1 | E01 | 049 | ACAGCTCA | TACTGCTC |
| 1 | E02 | 050 | GATCGAGT | GACGAACT |
| 1 | E03 | 051 | AGCGTGTT | CTTCGCAA |
| 1 | E04 | 052 | GTTACGCA | ATGGCGAT |
| 1 | E05 | 053 | TGAAGACG | ACATGCCA |
| 1 | E06 | 054 | ACTGAGGT | GTCAACAG |
| 1 | E07 | 055 | CGGTTGTT | GTGGTATG |
| 1 | E08 | 056 | GTTGTTCG | CCAACTTC |
| 1 | E09 | 057 | GAAGGAAG | GACGTCAT |
| 1 | E10 | 058 | AGCACTTC | ACGTCCAA |
| 1 | E11 | 059 | GTCATCGA | GATCCACT |
| 1 | E12 | 060 | TGTGACTG | AGCCTATC |
| 1 | F01 | 061 | CAACACCT | AGCTACCA |
| 1 | F02 | 062 | ATGCCTGT | AGATTGCG |
| 1 | F03 | 063 | CATGGCTA | CACACATC |
| 1 | F04 | 064 | GTGAAGTG | GAGCAATC |
| 1 | F05 | 065 | CGTTGCAA | ATAGAGCG |
| 1 | F06 | 066 | ATCCGGTA | GACCGATA |
| 1 | F07 | 067 | GCGTCATT | CAGACGTT |
| 1 | F08 | 068 | GCACAACT | CTGAACGT |
| 1 | F09 | 069 | GATTACCG | TTGGACTG |
| 1 | F10 | 070 | ACCACGAT | GTCTGCAA |
| 1 | F11 | 071 | GTCGAAGA | CCACATTG |
| 1 | F12 | 072 | CCTTGATC | GATGGAGT |
| 1 | G01 | 073 | AAGCACTG | AGGTCAAC |
| 1 | G02 | 074 | TTCGTTGG | TACACACG |
| 1 | G03 | 075 | TCGCTGTT | CAAGTCGT |
| 1 | G04 | 076 | GAATCCGA | AGCTAGTG |
| 1 | G05 | 077 | GTGCCATA | CTCCTAGT |
| 1 | G06 | 078 | CTTAGGAC | ACTCCTAC |
| 1 | G07 | 079 | AACTGAGC | CAATCAGG |
| 1 | G08 | 080 | GACGATCT | TCGTGCAT |
| 1 | G09 | 081 | ATCCAGAG | TAACGTCG |
| 1 | G10 | 082 | AGAGTAGC | AAGGCGTA |
| 1 | G11 | 083 | TGGACTCT | TCTTACGG |
| 1 | G12 | 084 | TACGCTAC | CGTGTGAT |
| 1 | H01 | 085 | GCTATCCT | AACAGGTG |
| 1 | H02 | 086 | GCAAGATC | AGTCGAAG |
| 1 | H03 | 087 | ATCGATCG | TGGAAGCA |
| 1 | H04 | 088 | CGGCTAAT | CTCGTTCT |
| 1 | H05 | 089 | ACGGAACA | ACGAGAAC |
| 1 | H06 | 090 | CGCATGAT | AAGCCTGA |
| 1 | H07 | 091 | TTCCAAGG | CTACAAGG |
| 1 | H08 | 092 | CTTGTCGA | CGATGTTC |
| 1 | H09 | 093 | GAGACGAT | ACCGGTTA |
| 1 | H10 | 094 | TGAGCTAG | GAACGGTT |
| 1 | H11 | 095 | ACTCTCGA | CTGTACCA |
| 1 | H12 | 096 | CTGATCGT | GCGCATAT |

**S2 Table.** Contd.

| Plate | Well | Index_ Pair_ID | i5_Sequence | i7_Sequence | Plate | Well | Index_ Pair_ID | i5_Sequence | i7_Sequence |
|---|---|---|---|---|---|---|---|---|---|
| 2 | A01 | 097 | CGACCATT | TGATAGGC | 2 | E01 | 145 | AGTGCAGT | ATTCCGCT |
| 2 | A02 | 098 | GATAGCGA | CATCCAAG | 2 | E02 | 146 | TTGATCCG | AAGCTCAC |
| 2 | A03 | 099 | AATGGACG | GTGAGACT | 2 | E03 | 147 | TGCCATTC | TGATCACG |
| 2 | A04 | 100 | CGCTAGTA | CTGATGAG | 2 | E04 | 148 | CTTGCTGT | CAATGCGA |
| 2 | A05 | 101 | TCTCTAGG | ACGGTACA | 2 | E05 | 149 | CCTACTGA | ATGCGTCA |
| 2 | A06 | 102 | ACATTGCG | CTCGACTT | 2 | E06 | 150 | CCAAGTTG | TACATCGG |
| 2 | A07 | 103 | TGAGGTGT | ACAACGTG | 2 | E07 | 151 | TGATCGGA | ACTGCGAA |
| 2 | A08 | 104 | AATGCCTC | TGCTGTGA | 2 | E08 | 152 | TAGTTGCG | TCTGTCGT |
| 2 | A09 | 105 | CTGGAGTA | CCAAGTAG | 2 | E09 | 153 | GTCTGATC | CTCAAGCT |
| 2 | A10 | 106 | GTATGCTG | AACTGAGG | 2 | E10 | 154 | CGTTATGC | AACCACTC |
| 2 | A11 | 107 | TGGAGAGT | AGGTAGGA | 2 | E11 | 155 | GCTCTGTA | CTTACAGC |
| 2 | A12 | 108 | CGATAGAG | TTCGCCAT | 2 | E12 | 156 | TTACCGAG | AGTCTTGG |
| 2 | B01 | 109 | CTCATTGC | CAGGTAAG | 2 | F01 | 157 | GCCATAAC | CACGCAAT |
| 2 | B02 | 110 | ACCAGCTT | GTATCGAG | 2 | F02 | 158 | CTCAGAGT | AGCTTCAG |
| 2 | B03 | 111 | GAATCGTG | TTCACGGA | 2 | F03 | 159 | CGAGACTA | CCTCGTTA |
| 2 | B04 | 112 | AGGCTTCT | GAGCTCTA | 2 | F04 | 160 | TGTGCGTT | TGAGACGA |
| 2 | B05 | 113 | CAGTTCTG | GTCAGTCA | 2 | F05 | 161 | TTCAGGAG | CACAGGAA |
| 2 | B06 | 114 | TTGGTGAG | CACGTCTA | 2 | F06 | 162 | GACTATGC | ACTCAACG |
| 2 | B07 | 115 | CATTCGGT | AATTCCGG | 2 | F07 | 163 | AGGTTCGA | AAGCGACT |
| 2 | B08 | 116 | TGTGAAGC | TCTAGGAG | 2 | F08 | 164 | AGTCTGTG | CCTACCTA |
| 2 | B09 | 117 | TAAGTGGC | ATCCGTTG | 2 | F09 | 165 | ACCTAAGG | ATCTCCTG |
| 2 | B10 | 118 | ACGTGATG | GATAGCCA | 2 | F10 | 166 | TGCAGGTA | TCACGATG |
| 2 | B11 | 119 | GTAGAGCA | TATGACCG | 2 | F11 | 167 | AAGGACAC | CCACAACA |
| 2 | B12 | 120 | GTCAGTTG | CGATTGGA | 2 | F12 | 168 | CAACCTAG | AGGTCTGT |
| 2 | C01 | 121 | ATTCGAGG | ACAAGCTC | 2 | G01 | 169 | CTGACACA | AGAAGGAC |
| 2 | C02 | 122 | GATACTGG | GAACCTTC | 2 | G02 | 170 | ACTCGTTG | GCGTATCA |
| 2 | C03 | 123 | GCCTTGTT | AGCGAGAT | 2 | G03 | 171 | AGCTCCTA | CAACACAG |
| 2 | C04 | 124 | TTGGTCTC | CCGTAACT | 2 | G04 | 172 | TACATCGG | TCCACGTT |
| 2 | C05 | 125 | CCGACTAT | TCAGACAC | 2 | G05 | 173 | CACAAGTC | ATCGCAAC |
| 2 | C06 | 126 | GTCCTAAG | CGAAGTCA | 2 | G06 | 174 | CGGATTGA | ACGTCGTT |
| 2 | C07 | 127 | ACCAATGC | GTGATCCA | 2 | G07 | 175 | AGTCGACA | CGAATACG |
| 2 | C08 | 128 | GATGCACT | ACTGGTGT | 2 | G08 | 176 | GTCTCCTT | TGCTTGCT |
| 2 | C09 | 129 | GCTGGATT | CTAACCTG | 2 | G09 | 177 | GAGATACG | CTCGAACA |
| 2 | C10 | 130 | ATGGTTGC | AGCCAACT | 2 | G10 | 178 | ATCGGTGT | ACATGGAG |
| 2 | C11 | 131 | CAGAATCG | CCAGTTGA | 2 | G11 | 179 | TCTCGCAA | ACAAGACG |
| 2 | C12 | 132 | GAACGCTT | AAGTGCAG | 2 | G12 | 180 | TCTAACGC | CGCCTTAT |
| 2 | D01 | 133 | TCGAACCA | AACCGTGT | 2 | H01 | 181 | CAATCGAC | AGCAGACA |
| 2 | D02 | 134 | CTATCGCA | CGCGTATT | 2 | H02 | 182 | GAGGACTT | GTTAAGCG |
| 2 | D03 | 135 | TACGGTTG | AGTTCGCA | 2 | H03 | 183 | TGGAGTTG | CATGGATC |
| 2 | D04 | 136 | GAGATGTC | TAGTCAGC | 2 | H04 | 184 | CTAGGCAT | ACAGAGGT |
| 2 | D05 | 137 | CTTACAGC | AACACCAC | 2 | H05 | 185 | CTCTACTC | TAAGTGGC |
| 2 | D06 | 138 | AGGAGGAA | GTAAGCAC | 2 | H06 | 186 | AGAAGCGT | AGTCAGGT |
| 2 | D07 | 139 | GACGAATG | GTCCTTGA | 2 | H07 | 187 | TCGAAGGT | GCCTTAAC |
| 2 | D08 | 140 | GAAGAGGT | CAGGTTCA | 2 | H08 | 188 | GTCGGTAA | GTTGGCAT |
| 2 | D09 | 141 | CGTCAATG | CCAACACT | 2 | H09 | 189 | ACGATGAC | CAACCTCT |
| 2 | D10 | 142 | TACCAGGA | GAGAGTAC | 2 | H10 | 190 | TCCGTATG | TGGATGGT |
| 2 | D11 | 143 | CGTACGAA | AGATACGG | 2 | H11 | 191 | CTAGGTGA | CTATCCAC |
| 2 | D12 | 144 | GACTTAGG | GTTCTTCG | 2 | H12 | 192 | CATTGCCT | GATCTCAG |

**S2 Table.** Contd.

| Plate | Well | Index_ Pair_ID | i5_Sequence | i7_Sequence |
|---|---|---|---|---|
| 3 | A01 | 193 | ACCTTCTC | GAACGAAG |
| 3 | A02 | 194 | TCGTGGAT | ACCTAGAC |
| 3 | A03 | 195 | GTTCATGG | TACGACGT |
| 3 | A04 | 196 | TAGGATGC | TTGAGCTC |
| 3 | A05 | 197 | CATGGAAC | AGTACACG |
| 3 | A06 | 198 | GCTTAGCT | TGTCAGTG |
| 3 | A07 | 199 | CTAACTCG | GACTACGA |
| 3 | A08 | 200 | ACCATGTG | TTACGTGC |
| 3 | A09 | 201 | TCAGACGA | ACTGCTTG |
| 3 | A10 | 202 | TATCAGCG | GCCTATGT |
| 3 | A11 | 203 | AGCAGATG | GTACCACA |
| 3 | A12 | 204 | AACGGTCA | TAGTGGTG |
| 3 | B01 | 205 | CGAACTGT | ATACGCAG |
| 3 | B02 | 206 | TCCGAGTT | AAGACCGT |
| 3 | B03 | 207 | TTCTCTCG | CTCCAATC |
| 3 | B04 | 208 | ATTCTGGC | TCTGGACA |
| 3 | B05 | 209 | ACTGCTAG | AACACTGG |
| 3 | B06 | 210 | CATAACGG | TTGGTGCA |
| 3 | B07 | 211 | CAGTCTTC | CCTGTCAA |
| 3 | B08 | 212 | TGCCTCTT | CTATGCCT |
| 3 | B09 | 213 | ACTGTGTC | TTCGGCTA |
| 3 | B10 | 214 | GTATTGGC | ACCGACAA |
| 3 | B11 | 215 | CGATGCTT | CGTAGATG |
| 3 | B12 | 216 | AAGGCTGA | CTGTATGC |
| 3 | C01 | 217 | AGTCAGGA | GTTGCTGT |
| 3 | C02 | 218 | CAGGTATC | AGAACCAG |
| 3 | C03 | 219 | TCTCCGAT | GATGTCGA |
| 3 | C04 | 220 | TTCAGCCT | AGGAGGTT |
| 3 | C05 | 221 | TCTGAGAG | AATCGCTG |
| 3 | C06 | 222 | TTAGGTCG | AGTGACCT |
| 3 | C07 | 223 | CTCTGGTT | CGAATTGC |
| 3 | C08 | 224 | GCGTTCTA | CAAGAAGC |
| 3 | C09 | 225 | TCACGTTC | CACCAGTT |
| 3 | C10 | 226 | AGGATGGT | GTATTCCG |
| 3 | C11 | 227 | GTGTTCCT | TTCGAAGC |
| 3 | C12 | 228 | GTAGCATC | AGACCTTG |
| 3 | D01 | 229 | AGGATCTG | CCAAGGTT |
| 3 | D02 | 230 | GACAAGAG | ACGTATGG |
| 3 | D03 | 231 | TTACGGCT | AAGGACCA |
| 3 | D04 | 232 | GCTGTTGT | TATGCGGT |
| 3 | D05 | 233 | AACCGAAG | AAGGAAGG |
| 3 | D06 | 234 | TCTGCTCT | AGCGTGTA |
| 3 | D07 | 235 | CTCAGCTA | TCTACGCA |
| 3 | D08 | 236 | CTTCACCA | TGGCTCTT |
| 3 | D09 | 237 | GATCGTAC | CCTTCCAT |
| 3 | D10 | 238 | CTACAGTG | ATACTGGC |
| 3 | D11 | 239 | TCGAGTGA | AACCTACG |
| 3 | D12 | 240 | CAAGTGCA | CATACTCG |

| Plate | Well | Index_ Pair_ID | i5_Sequence | i7_Sequence |
|---|---|---|---|---|
| 3 | E01 | 241 | CGAGTATG | TGCACTTG |
| 3 | E02 | 242 | CGTAGGTT | TCACTCGA |
| 3 | E03 | 243 | GCCAGTAT | CACTGTAG |
| 3 | E04 | 244 | ATGGAAGG | GTACGATC |
| 3 | E05 | 245 | AAGAGCCA | TGGTGAAG |
| 3 | E06 | 246 | TGCGTAGA | TAGCTGAG |
| 3 | E07 | 247 | TACACGCT | AGAGCAGA |
| 3 | E08 | 248 | CCTTCCTT | CTTCGGTT |
| 3 | E09 | 249 | ACCGCATA | ACAACAGC |
| 3 | E10 | 250 | TGGTCCTT | AGCCGTAA |
| 3 | E11 | 251 | CCATACGT | CTCTTGTC |
| 3 | E12 | 252 | AACCTTGG | CAGATCCT |
| 3 | F01 | 253 | CAAGGTCT | GATGCTAC |
| 3 | F02 | 254 | GCTTCGAA | AGGAACAC |
| 3 | F03 | 255 | CGGAATAC | ACCATCCT |
| 3 | F04 | 256 | AACTGGTG | GAACGTGA |
| 3 | F05 | 257 | GCTTCTTG | TAGAACGC |
| 3 | F06 | 258 | GCAATTCG | AACCAGAG |
| 3 | F07 | 259 | AGGTCACT | CGACCTAA |
| 3 | F08 | 260 | CAGCGATT | CTCTCAGA |
| 3 | F09 | 261 | AACCTCCT | AGGCTGAA |
| 3 | F10 | 262 | TCGACATC | ATCGGAGA |
| 3 | F11 | 263 | CTGGTTCT | GATACCTG |
| 3 | F12 | 264 | ACAGCAAC | TCCTGACT |
| 3 | G01 | 265 | GCATACAG | TCAGCCTT |
| 3 | G02 | 266 | CATCTACG | AAGCATCG |
| 3 | G03 | 267 | TTGTCGGT | GCCAATAC |
| 3 | G04 | 268 | TAGCCGAA | GACACAGT |
| 3 | G05 | 269 | AGGCATAG | AAGAGGCA |
| 3 | G06 | 270 | TTGACAGG | GAAGACTG |
| 3 | G07 | 271 | TGCACCAA | CCGTTATG |
| 3 | G08 | 272 | CCAGTGTT | CTAGCAGT |
| 3 | G09 | 273 | TGTCCAGA | GCCAGAAT |
| 3 | G10 | 274 | GATTGGAG | CGAGAGAA |
| 3 | G11 | 275 | ACGGTCTT | AACTCGGA |
| 3 | G12 | 276 | CTGCGTAT | ACAGTTCG |
| 3 | H01 | 277 | CACCACTA | TGACCGTT |
| 3 | H02 | 278 | TGTGGTAC | CATCTGCT |
| 3 | H03 | 279 | ACATAGGC | CGCTGATA |
| 3 | H04 | 280 | CAAGCAGT | TCGTCTGA |
| 3 | H05 | 281 | GCACGTAA | CACATGGT |
| 3 | H06 | 282 | TCGTAGTC | CGAGTTAG |
| 3 | H07 | 283 | CACTGACA | AGCTAAGC |
| 3 | H08 | 284 | CGTGTACT | GTTCCATG |
| 3 | H09 | 285 | GAGCTCAA | GCATCCTA |
| 3 | H10 | 286 | ACGTCGTA | CCATGAAC |
| 3 | H11 | 287 | GTCTAGGT | ATCCACGA |
| 3 | H12 | 288 | CTTCGTTC | GAGAAGGT |
