## Supplementary material for "A genomewide association study for bristle number variation in *Drosophila melanogaster*": S2_Text.pdf

**S2 Text.** Ingredients and brief protocol for Macdonald lab apple juice agar plates.

<u>Notes</u>

- Protocol makes ~1-liter of media which is sufficient for ~20 large (100-mm) petri dishes
- Agar (cat num: 66-111) and tegosept (cat num: 20-258) are from Genesee Scientific
- Apple juice and cane sugar are store bought

1) Mix the following in a 2-liter glass beaker: 750-ml $dH_2O$
20-g Agar

2) Add stir bar, cover beaker with saran wrap.

3) Stir / heat the mix at 380°C and 280-310 rpm for ~20 min, or until it vigorously boils.
*NOTE: The mix will not go clear. It will also be thick, and can easily burn, so want rpm of the stir bar to be high, and/or frequently stir with a spoon.*

4) Reduce heat to 325°C, remove beaker from the hotplate, and while stirring with spoon, add the following to the mixture: 250-ml Apple juice
25-g Cane sugar

5) Once hotplate is at 325°C, return beaker to hotplate, and stir at 320-rpm for 15 min.
*NOTE: Will not start to really boil until near the end of this time period; this is OK.*

6) Move the mixture to an orbital shaker, and cool to ~60°C (~30-min)
*NOTE: Avoid vigorous shaking to limit bubbles.*

7) Make up the following, and add to the cooling mixture at ~70°C: 1-g Tegosept
Make up 5-ml with 95% Ethanol
Shake to dissolve tegosept

8) Pour petri dishes.
*NOTE 1: Generally want dishes only ~1/2 full.*
*NOTE 2: Bubbles in the plates cause problems for egg collection; Avoid getting them, but if you do, remove by flaming agar surface with a bunsen burner.*
*NOTE 3: Immediately you are done with the glass cooking beaker, place in sink and fill with hot water; this will dramatically simplify cleaning.*

9) Wrap stacks of plates in saran wrap and store at 4°C.
