## Supplementary material for "A genomewide association study for bristle number variation in *Drosophila melanogaster*": S3_Text.pdf

**S3 Text.** Macdonald lab egg collection protocol.

Materials needed:

- 1X PBS - Diluted from 10X PBS (Invitrogen, AM9624) with $dH_2O$
- Live yeast paste - Mix $dH_2O$ 1:1 with active dry yeast granules (Genesee Scientific, 62-103) and mix until the consistency of toothpaste is achieved
- Apple juice agar plates (See Supplementary Text S2)
- A 200-µl pipette tip with the tip removed such that the bore/hole is ~1/16th of an inch in diameter

Getting eggs:

- Put a small amount of yeast paste in the center of a series of 100-mm apple juice agar plates
- Place plates in population cage for several hours
  *NOTE 1: To get very large numbers of eggs provide the cage with large amounts of yeast paste (e.g., in a plastic weighing boat) for 24 hours before egg collection.*
  *NOTE 2: The longer the apple juice plates are present in the cage the more likely it is that first instar larvae will be collected along with eggs. This is generally not a problem, but could be for some applications.*

Collecting eggs:

- Remove apple juice plates from cage
- Use a thin spatula to remove any remaining yeast paste, along with any dead flies/parts, from the agar surface
- Pour a layer of 1X PBS onto the plate surface
- Use a paintbrush to gently dislodge eggs from the surface
- Pour the PBS-resuspended eggs into a 50-ml conical tube via a funnel
- Spray the plate surface with 1X PBS over the funnel to ensure all eggs are recovered
- Wait for the eggs to sink (2-3 minutes), then pour off as much of the 1X PBS as possible without losing eggs
- Add new 1X PBS to 50-ml tube to wash the eggs, wait for them to sink, discard the PBS, and repeat this entire process 3-4 times until the PBS is clear
- Remove all but ~5-ml of the PBS with a transfer pipette, making sure not to disturb the eggs
- Get a P200 pipettor with a wide bore tip, press the plunger down, slide the tip down the side of the egg "pellet" in the conical tube, then let go of the plunger to aggressively pull eggs/PBS into the tip
- Hold the pipettor so that the tip is 1/4-1/2 inch from the surface of the media in the destination vial/bottle, and aggressively press down on the plunger to fire the contents out onto the media
- Repeat for all necessary vials/bottles
  *NOTE: Use 12-µl eggs for standard narrow vials and 36-µl eggs for 6-oz bottles*
