## Supplementary material for "A genomewide association study for bristle number variation in *Drosophila melanogaster*": S4_Text.pdf

**S4 Text.** Macdonald lab 96-well plate DNA isolation protocol.

Items

| | |
|---|---|
| Equipment: | BioSpec MiniBeadBeater-96 (2,100 rpm shaking speed) |
| | 65°C 2-block dry bath (need 1 bath per 96-well plate) |
| | 37°C hybridization oven |
| | 20-200 µl 8-channel pipettor |
| | Eppendorf centrifuge 5810 with A-4-62-MTP plate rotor |
| Reagents: | Puregene Cell Kit (Qiagen, 158046) |
| | 100% isopropanol (= 2-propanol) |
| | 70% ethanol |
| | EB Buffer (Qiagen, 19086) |
| Consumables: | Round bottom 96-well 500-µl assay plates (Axygen, P-96-450R-C) |
| | Silicone sealing mats (Axygen, AM-2ML-RD) |
| | 0.2-ml Thermo-Fast non-skirted PCR plates (AB-0600, Thermo) |
| | Domed 8-cap strips (AB-0265, Thermo) |
| | MicroAmp clear adhesive films (4306311, Applied Biosystems) |
| | Glass 3.5 mm grinding balls (11079135, BioSpec) |
| | MicroAmp 96-well rubber mats (N8010550, Applied Biosystems) |

Protocol

1) Chill cell lysis solution on ice until it goes cloudy.

2) Place a round-bottom 96-well plate containing 1 fly and 1 grinding ball per well on ice.

3) Add 100-µl cold cell lysis solution to each well. Use kimwipe to dry top surface of plate to ensure a proper seal for the next step. Seal plate with silicone mat.

4) Using MiniBeadBeater-96, homogenize the flies for 45-sec. Spin down at 1,000 rpm for a few seconds. Replace silicone mat with adhesive film

5) Incubate plate in dry bath (65°C for 20-min.)
*NOTE: Remove blocks from dry bath. Put plate on base of bath, overlay with rubber sheet and with plexiglass sheet, then put the blocks on top.*

6) Remove plate from dry bath, and spin down at 1,000 rpm for a few seconds. Remove and discard seal. Use a kimwipe dry top surface of the plate. Allow plate to cool to RT.

7) Add 30-µl of the following dilute RNase A mix to each well. *Do not* touch the wells with the tips, and pipet out gently. No mixing is required.

| | 210 rxns | 410 rxns |
|---|---|---|
| $dH_2O$ (µl) | 6,195 | 12,095 |
| RNase A (µl) | 105 | 205 |

8) Use kimwipe to dry top surface of plate. Seal plate with adhesive film. Incubate plate in hybridization oven (37°C for 40-min.)
*NOTE: Plate should be raised off the base of the oven on an upturned pipet tip box lid.*

9) Remove plate from oven, and immediately remove and discard seal. Use a kimwipe dry top surface of the plate. Allow plate to cool to room temperature.
*NOTE: It is very important the plate reaches room temperature before moving to next step.*

10) Add 33-µl protein precipitation solution to each well. *Do not* touch the wells with the tips, and pipet out gently. Use kimwipe to dry top surface of plate. Seal plate with adhesive film.

11) Vortex plate on high speed for 15 seconds, and place on ice for 10-min (make sure the bottom surfaces of the wells are in contact with the ice.)

12) Spin plate in centrifuge at 4,000 rpm (2,755 × g) for 20-min.

13) Prior to the end of the centrifugation step [step 12], add 100-µl 100% isopropanol to each well of an ABgene 96-well PCR plate.

14) *Carefully* remove seal from centrifuged plate. Move 110-µl of the supernatent into the isopropanol-filled plate, and *gently* pipet up-and-down once to mix. Seal the plate with a rubber mat.
*NOTE: After mixing, do not blow out all the reagent mix from the tips as the isopropanol causes minor splashing.*

15) Spin plate in centrifuge at 4,000 rpm (2,755 × g) for 30-min.

16) *Carefully* remove plate/holder from centrifuge and remove the mat. Put a folded paper towel on the surface of the plate and invert plate/holder. Move the inverted plate/holder to another folded paper towel to dry surface, and repeat this once or twice. Turn plate/holder the right way up again.

17) Add 100-µl 70% ethanol to each well. Seal the plate with a rubber mat. Spin plate in centrifuge at 4,000 rpm (2,755 × g) for 8-min.

18) *Carefully* remove plate/holder from centrifuge and remove the mat. Put a folded paper towel on the surface of the plate and invert plate/holder. Move the inverted plate/holder to another folded paper towel, and spin inverted plate/holder/towel in centrifuge at 250 rpm for 1-2 seconds.

19) Dry plate for 20-30 minutes on bench at room temperature (plate should be right way up). Add 10-µl EB, making sure the solution falls to the bottom of the wells. Seal plate with strip caps, and resuspend DNA by leaving overnight at room temperature.

20) Mix the DNA by removing the plate from the holder and flicking it several times. Spin the plate at 3,000 rpm for 1-2 seconds.
