## Supplementary material for "A genomewide association study for bristle number variation in *Drosophila melanogaster*": S5_Text.pdf

**S5 Text.** Custom, in-house Tn5-based library preparation protocol.

Tagmentation Step

14.5 µl  $H_2O$
4 µl  TAPS-DMF Buffer
0.5 µl  Tn5 w/pre-annealed oligos (see Picelli et al. 2014 – PMID: 25079858)
1 µl  gDNA @ 8ng/µl
---------
20 µl  TOTAL

Thermocycle:  55ºC  7 min
10ºC  hold
*Place on ice as soon at hits 10ºC then immediately move to next step*

Kill Enzyme Step

Add to each sample with a repeater:
5 µl  0.2% SDS

Thermocycle:  55ºC  7 min
10ºC  hold

PCR Amplification

Add to each well of a new plate with a repeater:
4.0 µl  KAPA 2X MasterMix (Roche, KK2612)

Add to each well with a multichannel:
1.0 µl  i5 UDI @ 3.75 µM
1.0 µl  i7 UDI @ 3.75 µM
2.5 µl  killed, tagmented gDNA (from previous step)
---------
8.5 µl  TOTAL

Thermocycle:  72ºC  3 min
98ºC  2 min 45 sec
12 cycles of:
98ºC  15 sec
62ºC  30 sec
72ºC  1 min 30 sec
4ºC hold
