## Supplementary material for "A genomewide association study for bristle number variation in *Drosophila melanogaster*": S6_Text.pdf

**S6 Text.** Library bead-cleanup protocols.

<u>0.8X Bead Cleanup</u>
*Applied to 12 non-size selected pools in order to concentrate, remove small fragments (e.g. primer dimers), and switch buffer*

1) Add 80-µl of AMPure XP beads to 100-µl of sample
2) Mix, then incubate at room temperature for 5-min
3) Place on magnet for 5-min
4) Remove and discard supernatant
5) Wash with 80% EtOH
6) Repeat EtOH wash
7) Air dry on magnet for 5-min
8) Resuspend in 60-µl of 10mM Tris-HCl w/ 0.1% Tween-20 (pH 8.5)
9) Incubate at room temperature for 2-min
10) Place on magnet for 5-min
11) Move supernatent to new tube

<u>2X Bead Cleanup</u>
*Applied to 12 size-selected pools in order to concentrate and switch buffer*

1) Add 80-µl of AMPure XP beads to 40-µl of sample
2) Mix, then incubate at room temperature for 5-min
3) Place on magnet for 5-min
4) Remove and discard supernatant
5) Wash with 80% EtOH
6) Repeat EtOH wash
7) Air dry on magnet for 5-min
8) Resuspend in 20-µl of 10mM Tris-HCl w/ 0.1% Tween-20 (pH 8.5)
9) Incubate at room temperature for 2-min
10) Place on magnet for 5-min
11) Move supernatent to new tube
