## Supplementary material for "A genomewide association study for bristle number variation in *Drosophila melanogaster*": S8_Table.pdf

**S8 Table.** Set of bristle number candidate genes that are within 20-kb of a GWAS association hit (adjusted *P*-value < 0.05). Genes noted as "in cluster" are present within one of the 3 clusters of hits on 3L (ABN), X (SBN) and 2L (SBN).

| Pheno | Gene Symbol | Gene Name | FBgn | Gene Location | In Cluster |
|---|---|---|---|---|---|
| ABN | *sws* | swiss cheese | FBgn0003656 | X:7956820..7968236[-] | - |
| | *exd* | extradenticle | FBgn0000611 | X:15992468..15996509[+] | - |
| | *lola* | longitudinals lacking | FBgn0283521 | 2R:10481894..10543291[-] | - |
| | *psq* | pipsqueak | FBgn0263102 | 2R:10557888..10617280[+] | - |
| | *trn* | tartan | FBgn0010452 | 3L:13114271..13118077[+] | Yes (3L) |
| | *caps* | capricious | FBgn0023095 | 3L:13228692..13279642[+] | Yes (3L) |
| | *sens* | senseless | FBgn0002573 | 3L:13396228..13401125[-] | Yes (3L) |
| | *Abp1* | Actin binding protein 1 | FBgn0036372 | 3L:13483533..13486280[-] | Yes (3L) |
| | *DCTN1-p150* | Dynactin 1, p150 subunit | FBgn0001108 | 3L:13929387..13934656[+] | Yes (3L) |
| | *nuf* | nuclear fallout | FBgn0013718 | 3L:14190870..14233250[+] | Yes (3L) |
| | *fz* | frizzled | FBgn0001085 | 3L:14274343..14368639[+] | Yes (3L) |
| | *bbg* | big bang | FBgn0087007 | 3L:14412828..14536276[-] | Yes (3L) |
| | *Tollo* | Tollo | FBgn0029114 | 3L:15235619..15242832[+] | Yes (3L) |
| | *Root* | Rootletin | FBgn0039152 | 3R:24113286..24128193[+] | - |
| SBN | *tyn* | trynity | FBgn0284435 | X:140318..200663[+] | Yes (X) |
| | *G9a* | G9a | FBgn0040372 | X:245978..254650[+] | Yes (X) |
| | *y* | yellow | FBgn0004034 | X:356509..361245[+] | Yes (X) |
| | *ac* | achaete | FBgn0000022 | X:370031..370947[+] | Yes (X) |
| | *sc* | scute | FBgn0004170 | X:396060..397497[+] | Yes (X) |
| | *l(1)sc* | lethal of scute | FBgn0002561 | X:409721..410817[+] | Yes (X) |
| | *ase* | asense | FBgn0000137 | X:460747..463176[+] | Yes (X) |
| | *Appl* | beta amyloid protein precursor-like | FBgn0000108 | X:530501..579044[+] | Yes (X) |
| | *Dredd* | Death related ced-3/Nedd2-like caspase | FBgn0020381 | X:633622..635545[-] | Yes (X) |
| | *esg* | escargot | FBgn0287768 | 2L:15333866..15336138[+] | Yes (2L) |
| | *wor* | worniu | FBgn0001983 | 2L:15423297..15425580[-] | Yes (2L) |
| | *sna* | snail | FBgn0003448 | 2L:15476593..15478260[-] | Yes (2L) |
| | *CycE* | Cyclin E | FBgn0010382 | 2L:15727581..15748150[-] | Yes (2L) |
| | *Gli* | Gliotactin | FBgn0001987 | 2L:15756002..15762758[-] | Yes (2L) |
| | *Tektin-A* | Tektin A | FBgn0028902 | 2L:15854126..15856450[+] | Yes (2L) |
